# Discovery of 17 conserved structural RNAs in fungi

**DOI:** 10.1101/2021.02.01.429235

**Authors:** William Gao, Thomas A. Jones, Elena Rivas

## Abstract

Many non-coding RNAs with known functions are structurally conserved: their intramolecular secondary and tertiary interactions are maintained across evolutionary time. Consequently, the presence of conserved structure in multiple sequence alignments can be used to identify candidate functional non-coding RNAs. Here, we present a bioinformatics method that couples iterative homology search with covariation analysis to assess whether a genomic region has evidence of conserved RNA structure. We used this method to examine all unannotated regions of five well-studied fungal genomes *(Saccharomyces cerevisiae, Candida albicans, Neurospora crassa, Aspergillus fumigatus,* and *Schizosaccharomyces pombe*). We identified 17 novel structurally conserved non-coding RNA candidates, which include 4 H/ACA box small nucleolar RNAs, 4 intergenic RNAs, and 9 RNA structures located within the introns and untranslated regions (UTRs) of mRNAs. For the two structures in the 3′ UTRs of the metabolic genes *GLY1* and *MET13*, we performed experiments that provide evidence against them being eukaryotic riboswitches.

## 1 Introduction

Non-coding RNAs (ncRNAs) perform important roles in protein synthesis, transcriptional and translational gene regulation, catalysis, guiding protein-driven processes, scaffolding, and more. In many cases, these ncRNAs function through a structure that is conserved across many species, even as their primary sequence diverges over evolutionary time. A hallmark of conserved RNA structure is the presence of compensatory base pair substitutions, which are substitutions in two nucleotides that maintain Watson-Crick or wobble base pairing despite changes in nucleotide identity. The degree to which two positions display compensatory base pair substitutions across evolution can be quantified through the covariation (correlated variation) of the positions pertaining to these nucleotides in a multiple sequence alignment.

Though comparative genomics methods using covariation analysis have been used to study RNA structure [1, 2, 3, 4], many of these methods do not account for substitutions that may appear compensatory but actually just result from phylogenetic relationships between the sequences in the alignment [5]. To address these concerns, R-scape (RNA Structural Covariation Above Phylogenetic Expectation) couples covariation analysis with statistical testing to identify nucleotides that significantly covary beyond phylogenetic expectation [6, 7]. A statistically significant covariation suggests that two nucleotides form a base pair that is important to the function of the RNA because there has been selective pressure to maintain it through compensatory base pair substitutions.

Failure to identify conserved base pairs through covariation analysis can be due to two reasons. Since covariation, by definition, requires some variation between sequences, positions in an alignment may have no covariation because the sequences are too conserved to observe compensatory base pair substitutions. Alternatively, there may be enough variation in the alignment but still no evidence of conserved structure. To account for these two scenarios, R-scape reports not only the number of conserved base pairs detected but also covariation power – a measure of the variation in the alignment. Using R-scape covariation and power calculations for all pairs in an alignment, the method CaCoFold (Cascade covariation/variation Constrained Folding algorithm) proposes a structure which includes all base pairs that have statistical support and prohibits all base pairs that have power but show no covariation [8].

R-scape has been used to evaluate proposed structures of numerous known ncRNAs. For many structural ncRNAs in the RNA Families (Rfam) database which have annotated consensus structures [9], R-scape/CaCoFold have enhanced these often manually produced structural annotations: base pairs predicted by R-scape and incorporated into an RNA structure by CaCoFold are highly concordant with positions shown to be in contact in available crystal structures [8]. For some other identified ncRNAs, such as HOTAIR, ncSRA and Xist, R-scape provides evidence against evolutionarily conserved RNA structure [6].

These were examples of R-scape/CaCoFold being used to improve known structures or critically test proposed ones. Here, we take advantage of R-scape’s ability to evaluate statistical significance for a third task – discovery of new structural RNAs, by identifying regions with significant covariation. For this purpose, we deployed it to screen all unannotated regions in the nuclear genomes of five fungi with high-quality genome annotations: *Saccharomyces cerevisiae, Candida albicans, Neurospora crassa, Aspergillus fumigatus,* and *Schizosaccharomyces pombe.*

While screens for structural ncRNAs in fungi have been performed in the past, they have historically been dwarfed by the number of such screens in bacteria [10, 11, 12, 13, 14, 15]. A major reason is that in earlier years the dearth of available whole genome sequences of fungi undermined the comparative genomics analysis that many structural RNA gene finders rely on. Additionally, fungal genomes, though comparatively small among eukaryotic ones, are nonetheless larger than bacterial genomes. Consequently, the low specificity of previous methods made screens of fungal and larger eukaryotic genomes less reliable [16].

Still, many researchers have sought to identify structural ncRNAs in fungal genomes, with most of these studies focusing on budding yeast, *S. cerevisiae*. Using probabilistic models called stochastic context-free grammars (SCFGs), Lowe et al. identified 22 C/D box small nucleolar RNAs (snoRNAs) [17]. By using QRNA [18], McCutcheon et al. discovered three H/ACA box snoRNAs and five RNAs of unknown function (RUFs) [19]. Kavanaugh et al. used thermodynamic approaches to find additional RUFs and structural elements in the *5′* and *3′* untranslated regions (UTRs) of protein-coding genes [20]. As recently as 2016, Hooks et al. reconciled the differences in sensitivity and specificity of existing RNA gene finding tools by screening all 306 budding yeast spliceosomal introns with three RNA structure prediction algorithms, EvoFold [21], RNAz [22], and CMfinder [23], identifying 14 structural ncRNA candidates that were in consensus among all three programs [24].

While less studied than *S. cerevisiae* and its subphylum Saccharomycotina (see Fig. 1 for phylogenetic tree), other fungi, particularly those within the subphylum Pezizomycotina, have also been subject to genome-wide screens for structural ncRNAs. Using an experimental screen for small RNAs in the pathogenic *A. fumigatus,* Jöchl et al. discovered 30 ncRNAs, many of which have subsequently been shown to be structurally conserved in the genus *Aspergillus* and, in some cases, across Pezizomycotina [25]. Using a bioinformatics method involving CMfinder, Li and Breaker recently discovered 15 novel structural ncRNA candidates, with most found in Pezizomycotina but some also in Saccharomycotina [26].

**Figure 1:**
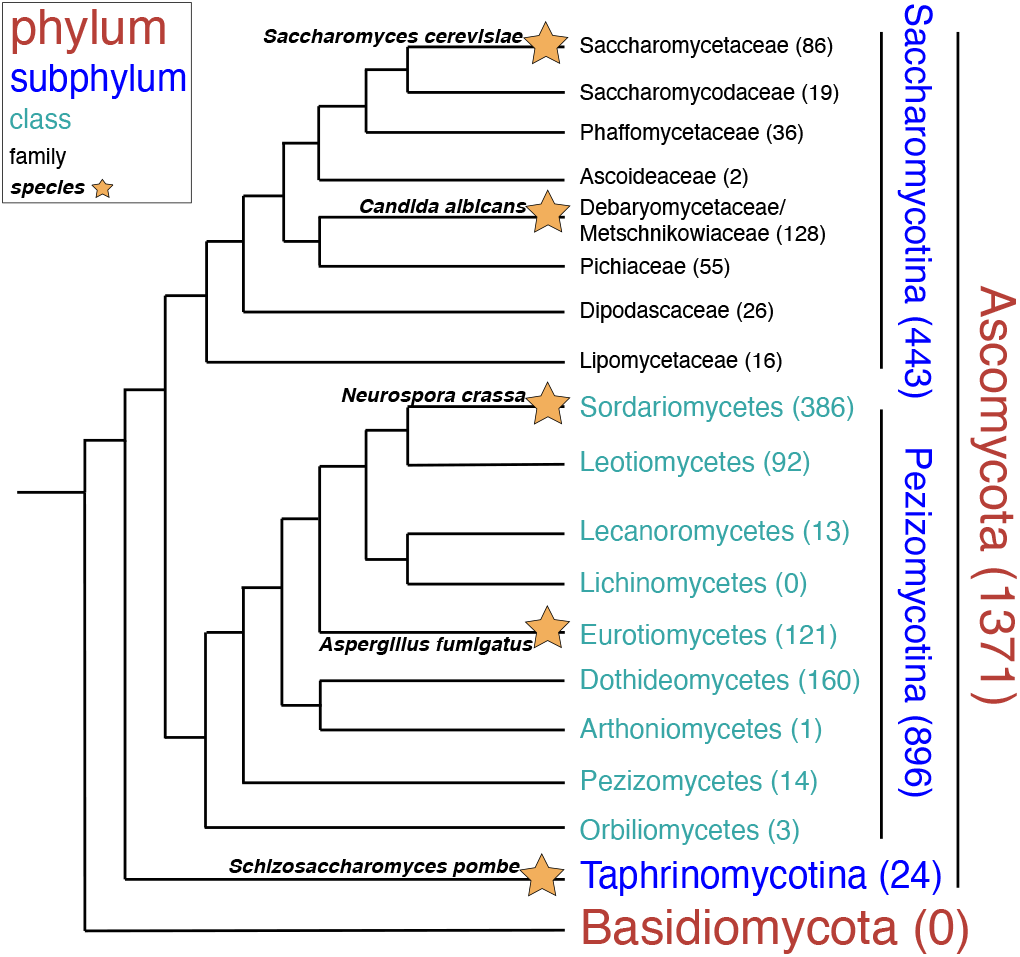
Phylogenetic distribution of the 1371 fungal genomes in the database. These genomes are all within the fungal phylum Ascomycota, which has been much more well-studied than the other major fungal phylum Basidiomycota. Ascomycota has three subphyla: Saccharomycotina, Pezizomycotina, and Taphrinomycotina. The taxonomy of all 1371 species was determined using the NCBI Taxonomy Browser. Per the legend, the font size and color reflect the taxonomic rank of each group listed in the phylogenetic tree. The number of genomes in each clade is listed after its name. In some cases, the sum of the number of genomes in lower levels does not equal the total number of genomes in a higher taxonomic level. This is because some genomes have been classified as *“incertae sedis,”* or of unknown taxonomic placement.

Overall, these screens have yielded many structural ncRNA candidates, some of which have since been validated experimentally. However, these previous screens utilized software programs that could result in false positives. This is because some of these structural RNA gene finders use both primary sequence conservation and consistency with a structure (i.e., compensatory base pair substitutions) as well as covariation when predicting structural ncRNAs. But of the three, structural conservation is the only evolutionary signature that distinguishes structural ncRNAs from other functional genomic features. For example, strong sequence conservation is also observed in many protein-coding sequences and some non-coding DNA sequences that bind to proteins, and a consistent RNA structure can be found in most genomic/DNA sequences, even random ones.

Importantly, not all covariation is due to RNA structure. Some of the previous software programs do not measure statistical significance of predicted base pairs, or do not consider the phylogenetic background of the alignment when calculating statistical significance. This likewise can result in a higher false positive rate by not accounting for observed covariation attributable to phylogeny. Because R-scape examines structural conservation alone through looking only at covariation and determines statistical significance of covarying positions by comparing against a null distribution that accounts for phylogenetic covariations, we control the false positive rate when using it to screen genomic regions for evidence of conserved structure.

Here, we have created a method to find novel conserved structural RNAs by using iterative homology search to build alignments that are then passed to R-scape for covariation analysis. Using this method in the genomes of five model organisms broadly distributed across the fungal kingdom, we have identified 17 novel conserved structural RNAs.

## 2 MATERIALS AND METHODS

### 2.1 Genomes and annotations

All alignments were constructed by searching against a database containing the most complete genomes available for 1371 fungi. These genomes were retrieved on May 19, 2019 from NCBI Genbank and represent the complete set of genomes designated as “Reference” or “Representative” from the phylum Ascomycota available at that time. A list of these genomes is provided in the Supplemental Data. Fig. 1 shows a phylogenetic tree of these genomes grouped into higher taxo-nomic ranks, with the branching order of Saccharomycotina based on a study from Shen et al. that used 1233 genes [27] and the branching order of Pezizomycotina based on a study from Spatafora et al. that used 5 genes [28].

Intergenic, intronic, and UTR sequences from five fungal genomes were used as query sequences in our screens. These genomes (S. *cerevisiae, C. albicans, N. crassa, A. fumigatus,* and *S. pombe*) were chosen because they are model organisms that have relatively thorough genome annotations, which provide genomic coordinates for protein-coding genes, non-coding RNAs, and other features such as transposable elements and pseudogenes. The annotation for the *S. cerevisiae* S288C genome (version=R64-2-1) was obtained from the *Saccharomyces Genome Database* website [29]. The annotations for *C. albicans, N. crassa, A. fumigatus*, and *S. pombe* were obtained from the NCBI Assembly website and have the following genome build accessions: *C. albicans* (GCA_000182965.3), *N. crassa* (GCF_000182925.2), *A. fumigatus* (GCF_000002655.1), and *S. pombe* (GCA_000002945.2).

### 2.2 Computational resources

Our method incorporates several computational tools. Iterative homology search with nhmmer (HMMER v3.2.1) was used to generate multiple sequence alignments [30]. Easel miniapps were used to extract query sequences from genomes in FASTA format (esl-sfetch) and to extract noncoding regions within an alignment in the flanked mode (esl-alimanip, esl-alimask). The Easel library for sequence analysis is also included in the HMMER v3.2.1 package. The structural RNA homology method Infernal (v1.1.2) was used to compare our candidate structures against existing Rfam models (cmscan), build and calibrate a new covariance model (cmbuild, cmcalibrate), and perform structure-informed homology search (cmsearch) [31, 32]. In the initial screen of all genomic regions, R-scape (v1.2.3) was used to perform all covariation analysis [6] as that was the most recent version available at the start of the study. The --cyk option was used to evaluate alignments produced from primary sequence homology searches, which lack a structural annotation. Using the CYK folding algorithm [33], R-scape produces a maximal covariation structural annotation for an alignment using all statistically significant covariations (E-value: 0.05) as base pairing constraints. For the alignments determined by our criteria to contain structural ncRNA candidates, CaCoFold was used to determine a final consensus structure (R-scape v1.5.16, with options -s -fold). In this setting, R-scape first evaluates the covariation support and power for the proposed structure and then analyzes all other possible pairs. CaCoFold then builds a structure that incorporates all covarying pairs and excludes all negative pairs (pairs with power but no covariation) [7, 8]. To examine the protein-coding potential of candidate structures, we used RNAcode v0.3 with default settings [34].

RNA-sequencing (RNA-seq) read mapping for Fig. 7c-d was performed using Galaxy [35]. All other computations in this paper were run on the Cannon cluster supported by the FAS Division of Science, Research Computing Group at Harvard University.

### 2.3 Aggregated p-values

To calculate the expected number of false positives in the screen, we used Fisher’s method [36] to produce an aggregated p-value for each RNA alignment.

For (*p*_1_,..,*p*_*n*_) p-values assumed to be independent and randomly distributed, Fisher’s method calculates the aggregated quantity

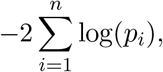

which follows a chi-squared distribution with 2*n* degrees of freedom. Thus, the aggregated p-value is given by

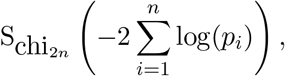

where S_chi_2*n*__ is the survival function of the chi-squared distribution with 2*n* degrees of freedom.

### 2.4 Experimental techniques

We experimentally tested if two of the identified conserved RNA structures were eukaryotic riboswitches. The *GLY1* 3′ UTR structure (163 nts, chr5:67409-67247) from *S. cerevisiae* and the *MET13* 3′ UTR structure (53 nts, scaffold4:71538-71590) from *Hyphopichia burtonii* were synthesized using in vitro transcription. As a positive control, we performed in-line probing and ITC experiments with the *xpt-pbuX* guanine riboswitch from *B. subtilis* (101 nts, 2320213-2320113). All sequences are provided in the Supplemental Data.

We performed two different assays to test whether these structures interacted with metabolites. The in-line probing assay was performed per the protocol by Regulski and Breaker [37]. Isothermal Titration Calorimetry (ITC) experiments were performed following the protocol by Gilbert and Batey [38], but with the MicroCal iTC200 calorimeter. Concentrations ranging from 1 to 30 *μ*M RNA were titrated against a corresponding 10-15 fold more concentrated ligand solution. A total volume of 40 *μ*L of ligand solution was injected at 30° C over 14-20 injections (each injection 2-3 *μ*L besides 0.5 *μ*L for the first injection) into the sample cell containing 200 *μ*L of RNA solution at a rate of 2 *μ*L/s, with 180 s intervals between injections, and a reference power of 10 *μ*cal/s.

Custom plasmids from Genscript were used as templates for the *GLY1* 3′ UTR structure and the guanine riboswitch. All oligonucleotides, including all primers used for PCR amplification along with the template for the *MET13* 3′ UTR structure were from Eton Bioscience. To verify their sequences, the RNAs used for in-line probing and ITC experiments were reverse-transcribed and sequenced with appropriate primers using the DNA sequencing service from Eton Bioscience. The metabolites tested with the *GLY1* 3′ UTR structure (glycine, L-threonine, L-serine, tetrahydrofolate, L-aspartate, guanine, adenine, pyridoxal phosphate, and thiamine pyrophosphate) were purchased from Sigma Aldrich. The same stock of guanine and adenine was also used with the guanine riboswitch. The candidate metabolites for the *MET13* 3′ UTR structure, SAM and SAH, were purchased from Cayman Chemical Company.

## 3 RESULTS

### 3.1 Method to identify conserved RNA structures

R-scape requires a multiple sequence alignment as input. To build these alignments, we performed iterative homology searches with nhmmer, a DNA/RNA homology finder [30]. This process involves building iterative probabilistic models called Hidden Markov Models (HMMs). Starting with a one query-sequence HMM, we search against a nucleic acid database of choice, accepting all homologs below a certain E-value threshold, and then building a new HMM from the generated alignment to use as the query for a subsequent search (Fig. 2). The false positive rate is specified by the E-value, which represents the number of sequences detected as homologous that one would expect by chance for the size of the database queried. Given a conservative E-value threshold between iterations, nhmmer can identify more distant homologs with each iterative search while maintaining a low false positive rate.

**Figure 2:**
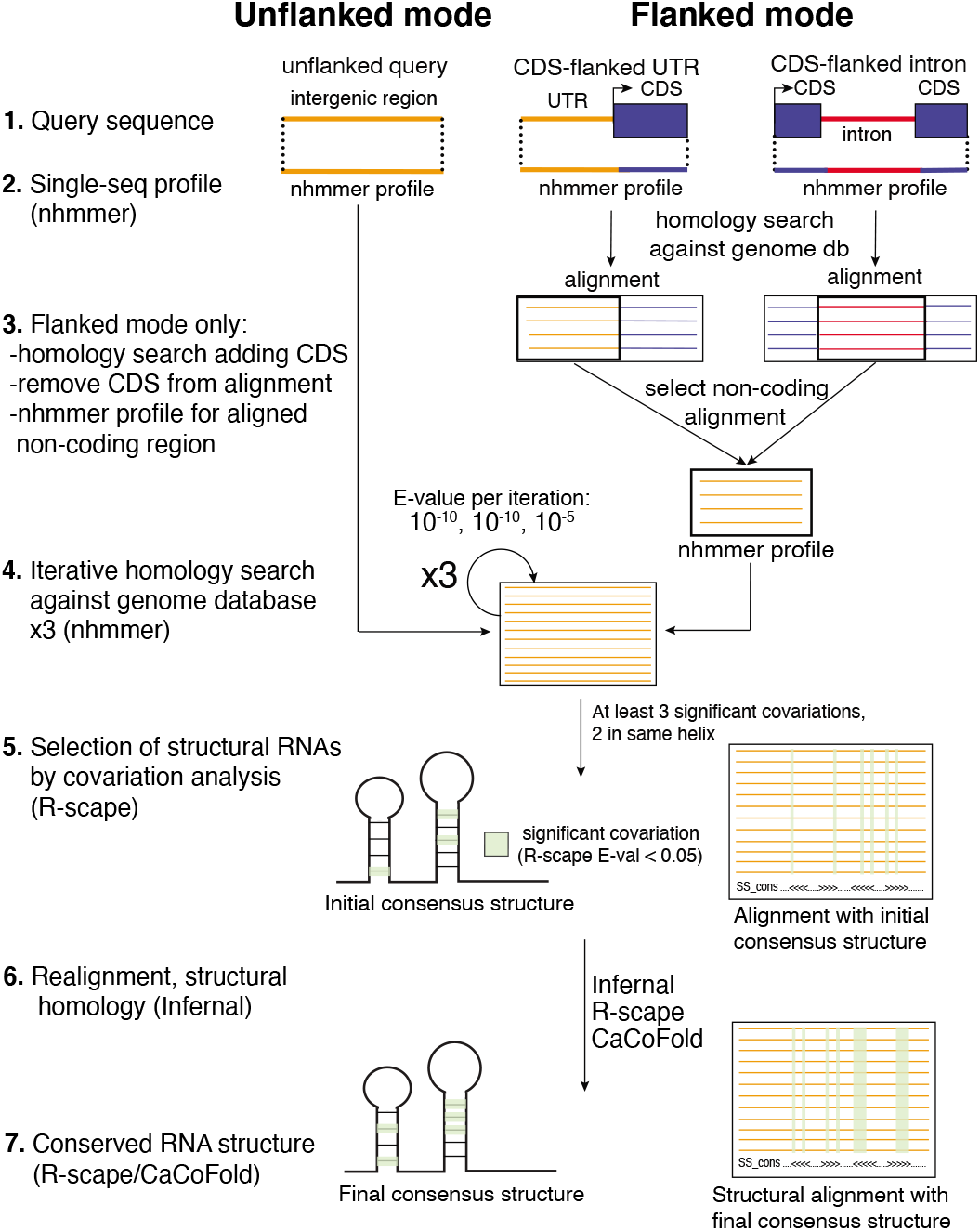
Description of the method: the flanked and unflanked modes. Our structural non-coding RNA finding method has two modes: an unflanked mode for finding standalone structural noncoding RNA genes and a coding sequence-flanked mode. In the flanked mode, sequences annotated as introns and UTRs are flanked by adjacent CDSes in the first homology search. Then, the noncoding portion of the produced alignment is extracted and all sequences with at least 50 nucleotides in this region are used to build an HMM that serves as the query for the subsequent homology search. We used the unflanked mode to screen all intergenic regions (IGRs) for standalone structural ncRNA genes and the flanked mode to screen all introns and UTRs for structures located in the non-coding regions of mRNAs.

#### Method optimization

Because covariation analysis is sensitive to the quality of the inputted alignment, we tested two parameters, the number of iterations and the E-value threshold, that could impact the covariation found in the final alignment. To devise the most informed set of parameters, we ran a series of different methods on a set of 80 known structural ncRNA genes which includes: 29 H/ACA box snoRNAs, 5S and 5.8S rRNAs, 39 tRNAs (one per anticodon), 5 spliceosomal RNAs, U3, Telomerase RNA, RNase MRP, RNase P, and SRP RNA. This positive set includes all annotated single-copy RNA genes, except for C/D box snoRNAs which tend to have very little covariation and power. Although they have numerous covariations, the small and large subunit rRNAs were excluded because they are not representative of the length and copy number of typical structural RNAs.

For each ncRNA, the input sequence from *S. cerevisiae* was flanked with intergenic sequence all the way to the upstream and downstream annotated features (Fig. S1). With this set of positive controls, we found that the method which yielded the most covariations without significantly diminishing returns involved three iterative homology searches, with the first two E-value thresholds at 10^-10^ and the third at 10^-5^. To avoid including possible RNA pseudogenes, which are no longer structurally constrained and thus weaken the covariation observed between conserved base pairs, we kept only the top scoring hit per genome.

#### The flanked and unflanked modes

While the above approach suffices for finding standalone ncRNA genes, which comprised our entire positive control set, we hypothesized that to detect homologs of ncRNA structures associated with protein-coding genes, it would be useful to rely on the adjacent coding sequences (CDSes) which are typically more conserved at the primary sequence level. Therefore, as a slight modification to the above “unflanked” mode, we also designed a flanked mode that exploits the higher primary sequence conservation of adjacent coding sequences to aid in detection of structured cis-regulatory elements (Fig. 2).

In the unflanked mode, the input sequence (an intergenic region) serves as the initial query for three rounds of primary sequence homology search. In the flanked mode, the input sequence (a non-coding region such as an intron or untranslated region flanked by coding regions) serves as the query for an initial round of homology search. The non-coding portion of this alignment is then extracted to serve as the query for three more rounds of homology search. The reason for removing the coding portion after the first step in the iteration is to avoid having the homology search focus only on the typically more-conserved coding region. Using the flanking mode was fundamental to identifying structures with low primary sequence conservation but significant covariation.

In both search modes, the final primary sequence-based alignment is passed to R-scape for covariation analysis. For structural ncRNA candidates, a covariance model of the alignment is then produced using Infernal and used for a structure-informed homology search to generate a final alignment. R-scape is then used on this alignment to identify covarying base pairs which CaCoFold uses to predict a structure.

#### Determination of query sequences

Having optimized our method, we next decided on which genomes to screen using it. Since our database contained 1371 fungal genomes within the kingdom Ascomycota, we wanted query genomes that span the breadth of this clade. Additionally, we sought to use genomes that are well-annotated. Starting with query genomes with high-quality annotations is important because this can greatly decrease the search space. In particular, it helps to avoid screening transposable elements, which can comprise a significant fraction of the genome, and because of their association with repetitive elements, can result in large numbers of conserved alignments. Additionally, high-quality annotations are essential for providing the genomic coordinates of protein-coding genes, and importantly, the boundaries of intronic sequences. UTRs, however, are rarely annotated, even for *S. cerevisiae.* Therefore, to use the flanked mode for UTRs, the entire intergenic region (IGR) adjacent to the most upstream or downstream annotated CDS of each mRNA was treated as the UTR and flanked either by the downstream CDS in the case of 5′ UTRs or by the upstream CDS in the case of 3′ UTRs (Fig. S2).

We also wanted the query genomes to be experimentally tractable organisms, which are typically the most well-annotated ones. With these three criteria in mind, we chose to query all unannotated regions of five fungal genomes: *S. cerevisiae, C. albicans, N. crassa, A. fumigatus*, and *S. pombe*. Initially, we also explored the possibility of doing a pan-fungal screen by including the available fungal genomes from Basidiomycota and other phyla. However, as demonstrated by how only two structures were found in more than one subphylum, it is unlikely that many structures would have been found in both phyla since they are even more divergent. TimeTree estimates the divergence time of the subphyla Saccharomycotina and Pezizomycotina as 590 million years ago (MYA) and that of the phyla Ascomycota and Basidiomycota as 723 MYA [39]. Moreover, no high-quality genome annotation was available from Basidiomycota at the time of our study.

For each of these five query genomes, we used the unflanked mode on each IGR and used the flanked mode on each intron and UTR sequence. The minimum IGR length screened was 100 nts, since anything shorter was empirically determined to have an E-value greater than 10^-10^ even for a perfect match. For IGRs longer than 1000 nts, windowing was performed with 500 nt overlaps. For example, a 2500 nt IGR would be divided into four windows: nts 1-1000, 501-1500, 1501-2500, and 2001-2500. For IGRs that were treated as UTRs for the flanked mode, the UTR was restricted to at most the 2000 nts closest to the adjacent CDS. The flanking CDS was also restricted to at most 1000 nts. For each query sequence, our method produces an alignment and structural covariation results from R-scape. Details of the input sequences and the alignments are given in Table S1.

#### Determination of conserved RNA structures

By examining the covariations observed in our positive control set, we devised the following criteria for classifying the alignment of an unannotated genomic region as containing a structural ncRNA candidate: (1) at least 3 total statistically significant (E-value: 0.05) covarying base pairs and (2) at least 2 statistically significant covarying base pairs located in the same stem within the proposed structure. Of the 80 positive control RNAs, 62 (78%) satisfy both criteria (Fig. S1). Additionally, the contents of each column of these positive controls were shuffled independently from each other to create new alignments that disrupt covariation but preserve the base composition and position-specific sequence conservation. 0% of these alignments, which served as negative controls, satisfied either criterion.

To allow for structure-informed homology search, we used Infernal to build covariance models of the candidates that satisfied these criteria [31, 32]. These covariance models were then used to search the whole genome database for additional homologs. Multiple sequences per genome were allowed in this alignment unlike before, although as shown in Table 1, most of the genomes only have one copy of each RNA. The structures we present in this paper are the ones proposed by CaCoFold for this final alignment.

**Table 1:**
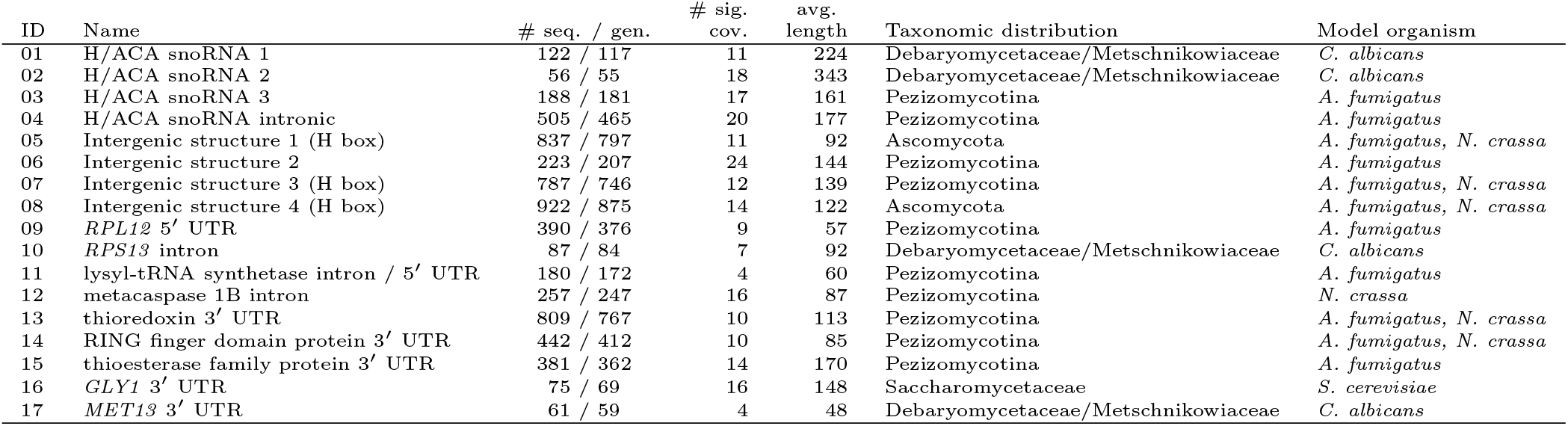
List of 17 conserved structural RNAs. In our screen of five fungal genomes, we discovered 4 H/ACA small nucleolar RNAs, 4 likely intergenic non-coding structural RNAs, and 9 non-coding structures associated with mRNAs. The structures are predominantly single copy per genome, as indicated by the similar number of sequences and genomes. The “taxonomic distribution” column is the taxonomic rank encompassing all the detected representatives (not all genomes in that taxonomic rank necessarily have the structure). Refer to Fig. 1 for more information of the taxonomy of the fungi included in our whole genome database. Additionally, the “model organism” column indicates which of the five query genomes have that structure. Additional information about location, coordinates and genomic context is given in Table S2.

Importantly, primary sequence homology search algorithms like nhmmer may not produce high quality alignments for covariation analysis since they find primary sequence homology rather than structural homology. Therefore, we also examined all genomic regions with fewer than 3 statistically significant base pairs in the final primary sequence alignment but at least one covarying base pair with an E-value less than 10^-4^, over 2 orders of magnitude smaller than the default E-value threshold of 0.05. In these cases, the structure proposed by R-scape from the primary sequence alignment was used to build a covariance model. This covariance model was then used to search the whole genome database. Covariation analysis was then performed on this final alignment and only candidates that satisfied the aforementioned two criteria (3 significant covariations, two of them in the same stem) were classified as candidate ncRNA structures. This additional step led to the detection of three candidate structures: conserved intergenic structure 4 (#08), the lysyl-tRNA synthetase intron / 5′ UTR structure (#11), and the *MET13* 3′ UTR structure (#17).

We removed 1528 regions that had more than three covariations between positions that were one or two nucleotides apart, suggesting that they may be interactions between positions of a codon within a protein-coding sequence. The average number of within-codon-like covariations across these alignments is 36, so they are likely unannotated protein-coding genes or pseudogenes. These covariations appear to be related to protein-coding evolution, and we are investigating them elsewhere.

To determine the novelty of all identified conserved RNA structures, we searched each of their representatives against all RNA structural families present in the Rfam database [9]. Overall, 106 regions were removed across the five query genomes: 35 rRNAs, 32 tRNAs, 25 snoRNAs, and 15 to other RNA genes (e.g. snRNAs, RNase P, Telomerase RNA). Many of the candidates that were removed, especially the ones in *S. cerevisiae*, are probably unannotated ncRNAs or pseudogenes (Table S3). For the remaining structures, there were either no significant hits (E-value: 0.05) to any families, or almost certainly spurious hits to Rfam models whose representatives are all in clades distant from fungi (e.g., a significant hit to a small RNA only found in *Drosophila*). All of these hits also had a score well below the score threshold specified for each Rfam model. Additionally, we conducted a literature search to determine if any of the structural ncRNA candidates were previously identified.

We also tested the protein-coding potential for the 17 structural RNA candidates using RNA-code [34]. Three candidates showed significant hits with p-values smaller than 0.05. Structure #02 has an p-value of 0.007 for a 12 nt open reading frame (ORF), Structure #05 has a p-value of 0.010 for a 21 nt ORF, and Structure #08 has a p-value of 0.027 for a 33 nt ORF. We believe that these are likely false positives due to the short predicted ORF length and relatively high p-values (most protein-coding multiple sequence alignments we ran with RNAcode had a hit with p-values smaller than 10^-16^, even for proteins under 100 amino acids). Additionally, as we discuss below, all three of these structures have features of H/ACA snoRNAs.

Our final list of 17 novel conserved structural RNAs is given in Table 1 and described in detail below. Information about location, coordinates and size of all RNAs is given in Table S2. A list with the 106 regions with an Rfam hit are given in Table S3. The final structural alignments of all 17 structural candidates as well as the nhmmer alignments after the three iterations are provided as Supplemental Data.

We have estimated the expected number of false positives for the whole fungi screen in which we tested a total of 134,000 alignments. We estimate the p-value threshold for detecting an RNA structure as the aggregated p-value of snR80, which is the RNA in the positive control that satisfies the selection criteria with the lowest amount of covariation. snR80 has three significant covariations of individual E-values of 6.5 ×10^-7^, 0.0053, 0.0204, which results in an aggregated p-value threshold of 2.1×10^-8^, according to Fisher’s method (see Methods). Thus, assuming all 134,000 alignments tested are not conserved structural RNAs, we should expect 0.003 total false positives for the whole screen.

### 3.2 Re-identification of recently discovered structures

In addition to the 17 novel structures presented here, our screen identified four other conserved RNA structures which are excluded from this final count since they have very recently been discovered by others. None have yet been included in the Rfam database. These are structures in the introns of *RPL7B* and *RPS13* identified by Hooks et al. [24], the *RPS2* 5′ UTR structure identified by Li and Breaker [26], and the *RPL9B* 3′ UTR structure identified by Gudipati et al. [40]. These structures were discovered using entirely different methods yet result in predicted structures nearly identical to ours, which underscores the soundness of our approach and theirs. Additionally, at least partly because our genome database contains many more genomes, we were able to extend the phylogenetic distribution of all these structures. The *RPL7B, RPS13,* and *RPL9B* structures are found in most of Saccharomycetaceae. We find the *RPS2* structure not only in Pezizomycotina, as reported by Li and Breaker, but also in over 300 Saccharomycotina genomes.

### 3.3 Discovery of four H/ACA box snoRNA candidates

Among the 17 novel conserved RNA structures, we identified four putative H/ACA snoRNAs. H/ACA snoRNAs guide the pseudouridylation of sites within rRNAs and other RNAs. These standalone ncRNA genes are so named because they all share an H box motif (ANANNA) and an ACA box motif (ACA) [41], with these motifs located at conserved distances from the pseudouridylation pockets [42]. In our screen of *C. albicans*, we identified two H/ACA snoRNAs in the Debaryomycetaceae / Metschnikowiaceae families within the subphylum Saccharomycotina (Fig. 3a-b).

**Figure 3:**
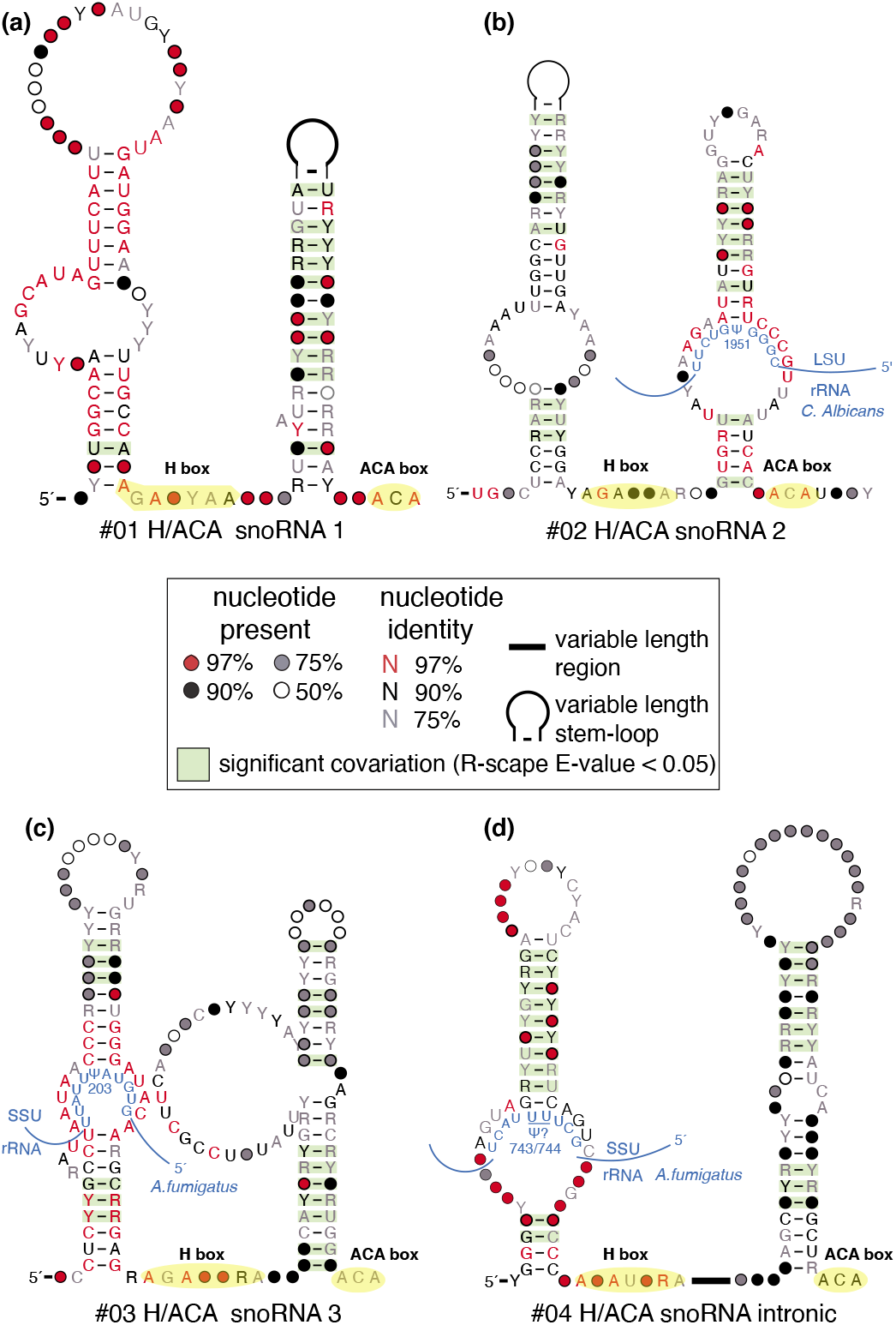
Four H/ACA snoRNA structural candidates. Three of these are intergenic **(a-c)** and the other **(d)** is consistently found in the intron of a vacuolar sorting protein. For each structure, the H box (ANANNA) and ACA box (ACA) motifs are highlighted in yellow and labeled. Nucleotides are colored according to degree of conservation per the legend. Base pairs highlighted in green are statistically significantly covarying. Putative interactions between the guide regions of the H/ACA snoRNAs and rRNAs are illustrated in blue, with the proposed site of pseudouridylation designated as ψ. R-scape uses R2R to draw the structures [43]. The sequences have been trimmed to highlight the region of covariation. Positions with more than 50% gaps are not displayed unless they are base paired.

In our screen of *A. fumigatus*, we found two additional H/ACA snoRNAs in Pezizomycotina. One of these is intergenic like the previous two (Fig. 3c) while the other consistently resides in the intron of a gene annotated as a “vacuolar sorting protein” by NCBI (Fig. 3d). To determine this, we examined the genomic context of the H/ACA snoRNA in the genome annotation of *A. fumigatus* as well as four other genomes with decent genome annotations: *Rasamsonia emersonii*, *Talaromyces stipitatus*, *Penicillium chrysogenum*, and *Ascodesmis nigricans*. Importantly, while the first four species are of the class Eurotiomycetes, *A. nigricans* is of the class Pezizomycetes. The estimated divergence time between *A. nigricans* and *T. stipitatus* is 442 million years ago (MYA) according to TimeTree [39], suggesting that the intronic location of this H/ACA snoRNA is highly conserved.

To guide the pseudouridylation of specific nucleotides on rRNAs, H/ACA snoRNAs often have highly conserved nucleotides in the interior loop of one or both of their hairpins. These nucleotides base pair with regions of the rRNA that are directly upstream and downstream of the modified uracil. For the second hairpin of H/ACA snoRNA 2 (#02), and the first hairpins of H/ACA snoRNAs 3 (#03) and H/ACA snoRNA intronic (#04), we used their high primary sequence conservation to identify plausible uracil targets within rRNAs. We used the consensus sequence of the snoRNA guide to identify a target on SSU (small subunit) and LSU (large subunit) rRNA sequences of the model organism containing each snoRNA. We searched for complementarity to the guide with at least 4 base pairs on both sides of the pseudouridylation site (must be a U), with G:U wobble base pairing allowed [42]. We found that LSU-1951 in *C. albicans* (corresponds to LSU-1955 in *S. cerevisiae*, although the snoRNA is not found there) is a putative target of H/ACA snoRNA 2’s second hairpin’s guide, SSU-203 in *A. fumigatus* (SSU-207 in *S. cerevisiae*) is a putative target of H/ACA snoRNA 3′s first hairpin, and SSU-743/744 in *A. fumigatus* (SSU-744/745 in *S. cerevisiae*) is a putative target of H/ACA snoRNA intronic’s first stem. We were unable to establish strong putative targets for the other hairpins due to lack of high primary sequence conservation. Consistent with their designation as novel H/ACA snoRNAs, we also found no previous computational or experimental determination of these Us as targets of any known snoRNAs [44, 45, 46].

As negative controls, we used 10000 shuffles of the target rRNA sequence to query for matches to the snoRNA guide with (1) as many base pairs and (2) fewer or equal number of wobble base pairs as the hypothesized target sequence. The false positive rates for the targets on snoRNAs #02, #03, and #04 were 0.017, 0.011, and 0.003, respectively.

### 3.4 Discovery of four conserved intergenic structural ncRNAs

Along with three of the four H/ACA snoRNA candidates, we also identified four other structural RNAs that are found in many different intergenic contexts across their representatives (Fig. 4). We refer to these structures as conserved intergenic structures 1-4. Intergenic structures 2 and 3 are found only in Pezizomycotina while intergenic structures 1 and 4 are also present in some of Saccharomycotina. These are the only two structures found in more than one subphylum. Intergenic structures 1,3, and 4 all have an H box motif and a single hairpin with a potential guide region, but lack a secondary hairpin and an ACA box. Thus, we have not listed them as H/ACA snoRNAs. Nonetheless, it may be possible that they perform similar functions as guide RNAs. While the H and ACA boxes both serve as nucleolar localization signals, previous research has shown that deletion of both is required to fully abolish localization [47]. Moreover, single-hairpin guide RNAs with an AGA box (rather than ACA) have been reported in several basal eukaryotic genomes and archaea [48, 49, 50, 51, 52]. Because of the strong conservation in their interior loops, we used the same approach as described above for the H/ACA snoRNA candidates to identify potential targets on rRNA for intergenic structures 1 and 4.

**Figure 4:**
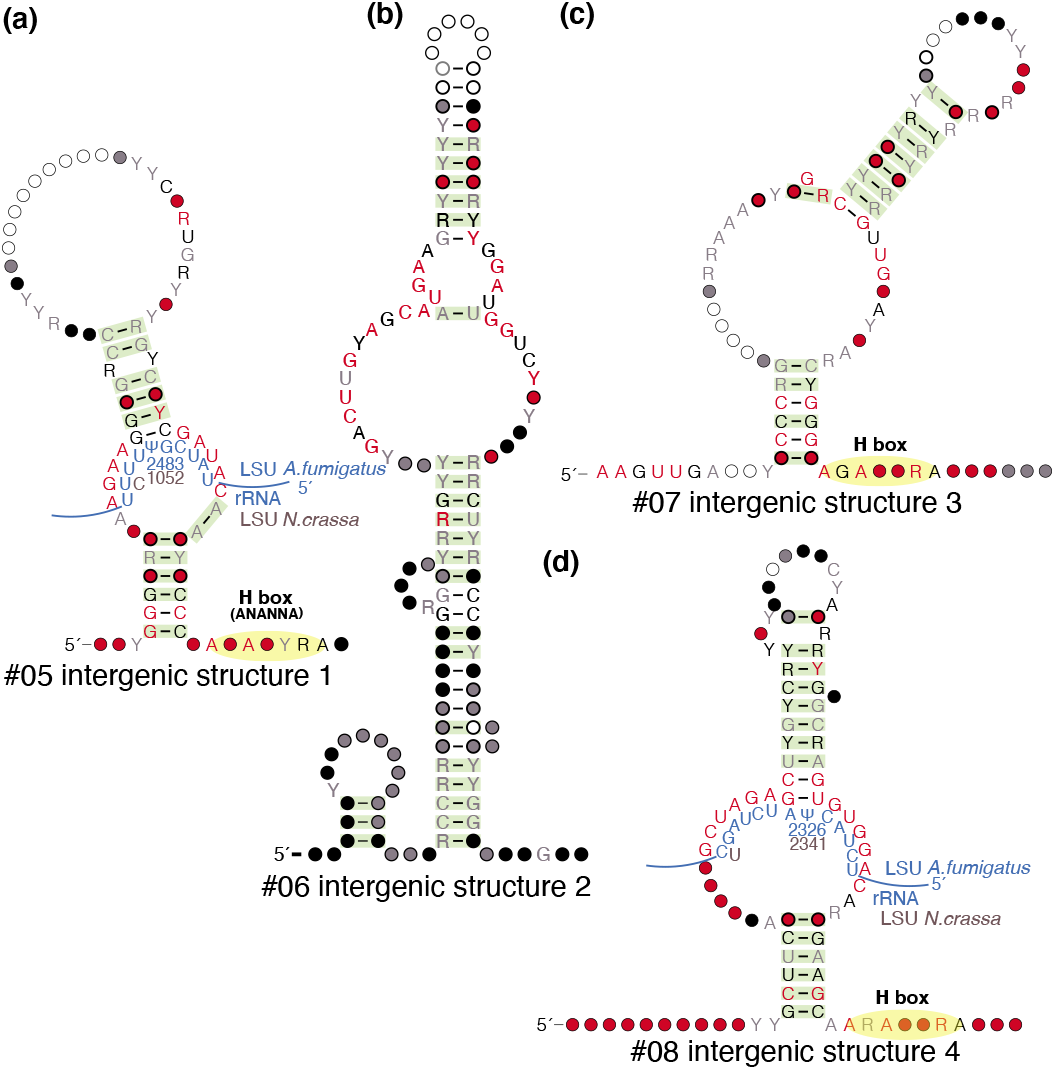
Four conserved intergenic structural non-coding RNA candidates. These intergenic structures are found in different genomic contexts across genomes with high quality annotations, so we hypothesize that they are standalone non-coding RNA genes. Intergenic structures 1, 3, and 4 have some but not all of the signatures associated to H/ACA snoRNAs. Refer to the legend from Fig. 3 for this and subsequent figures.

Intergenic structure 1 may target LSU-2483 in *A. fumigatus* (LSU-2504 in *S. cerevisiae*) and may target LSU-1052 in *N. crassa* (LSU-1039 in *S. cerevisiae*). Intergenic structure 4 may target LSU-2326 in *A. fumigatus* and LSU-1052 in *N. crassa* (LSU-2347 in *S. cerevisiae*). Intergenic structure 1, if it serves as a guide, appears to target two different sites in *A. fumigatus* and *N. crassa*, as has been shown for some snoRNAs [46, 17].

### 3.5 Discovery of nine ncRNA structures associated with mRNAs

We also identified nine RNA structural candidates located either in the introns or UTRs of mRNAs. Since the genomic coordinates of UTRs are typically not annotated, this determination was made by examining the genomic context of all representatives with available high-quality genome annotations (list provided in Supplemental Data), using a similar approach as described above with the intronic H/ACA snoRNA. In contrast to the eight standalone conserved intergenic structures discussed above, representatives of the nine structures described here are consistently adjacent to or within the same protein-coding gene across the different representative genomes. The direction of transcription of this adjacent gene was then used to assign the structure as located within either its 5′ or 3′ UTR.

We performed further targeted analysis of these 9 candidates associated with protein-coding genes. This involved creating a smaller database with each protein-coding gene and flanking sequences 3000 nts upstream and downstream. We used the Infernal covariance model of each structural candidate to search for homologs in this smaller database. This analysis increased the number of representatives for most of the candidates (the alignments are available in the Supplemental Data), but did not significantly change the phylogenetic distribution, except for structure #11, which went from 180 to 498 sequences, and structure #14, which went from 442 to 870. We still find that while most of the protein-coding sequences are detectable across all of Ascomycota, these non-coding structures are only present in a single subphylum or even smaller taxon.

#### Two ribosomal protein structural motifs

Many structural motifs associated with ribosomal proteins have been previously reported in bacteria and eukaryotes [53]. These structural motifs are typically involved in an autoregulatory feedback loop in which the ribosomal protein binds to its own pre-mRNA to regulate gene expression, often through regulation of splicing in eukaryotes [54]. Here, we report the discovery of two more ribosomal protein structural motifs, a 5′ UTR structure in *RPL12* (Fig. 5a) and an intronic structure in *RPS13* (Fig. 5b), found in our screens of *A. fumigatus* and *C. albicans,* respectively. Interestingly, Hooks et al. reported an *RPS13* intronic structure in *S. cerevisiae* and other species in the family Saccharomycetaceae [24]. We also found this structure but do not detect homology between it and the *C. albicans RPS13* structure using Infernal and fail to find any conserved structure in the intervening clades between Saccharomycetaceae and Debary-omycetaceae/Metschnikowiaceae (see Fig. 1 for phylogenetic tree). Thus, we currently present the Debaryomycetaceae/Metschnikowiaceae *RPS13* structure as distinct from the one previously discovered.

**Figure 5:**
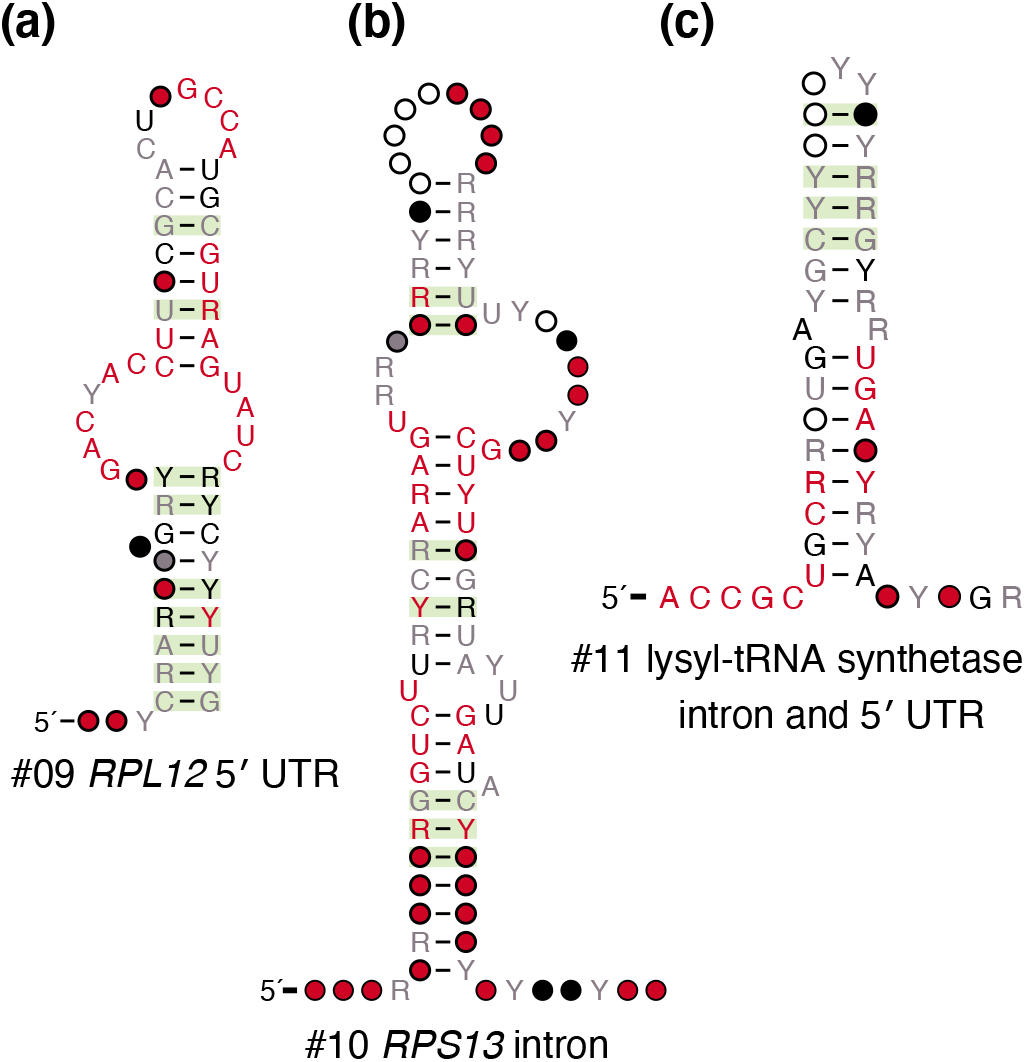
Three mRNA structures likely to engage in autoregulation. We discovered 3 non-coding RNA structures associated with protein-coding genes that are likely to interact with the proteins produced from their mRNAs to autoregulate their translation. These include: **(a)** a 5′ UTR structure located in the *RPL12* gene encoding a ribosomal protein, **(b)** a structure located in the sole intron of the *RPS13* gene which also encodes a ribosomal protein, and **(c)** a structure located in either the 5′ UTR or intron of a gene encoding lysyl-tRNA synthetase.

#### Lysyl-tRNA synthetase intronic and 5′ UTR structure

In our screen of *A. fumigatus*, we found a structure associated with the lysyl-tRNA synthetase mRNA, mostly in its first intron but also occasionally in the 5′ UTR, as seen in *Purpureocillium lilacinum* and *Neosartorya fischeri* (Fig. 5c).

As with ribosomal proteins, aminoacyl-tRNA synthetases have been shown to engage in autoregulation by directly binding to RNA motifs within their own mRNAs. For example, Frugier and Giegé showed that the aspartyl-tRNA synthetase in *S. cerevisiae* contains an additional N-terminal domain that is not involved in aminoacylation [55]. Instead, this domain binds a stretch of 87 nucleotides in its own mRNA’s 5′ UTR, with reporter constructs showing that this interaction results in reduced translation of the aminoacyl-tRNA synthetase. More recently, Levi and Arava have used RNA immunoprecipitation coupled with deep sequencing to show that most aminoacyl-tRNA synthetases in *S. cerevisiae* have a strong preference for their own mRNAs, with their own mRNA often being the top association [56]. Through experiments with the histidyl-tRNA synthetase, they identified a small structure in its mRNA that binds to the protein and represses translation. They propose the following model of autoregulation: the aminoacyl-tRNA synthetase has a tRNA primary target (high affinity), and a secondary target on its own mRNA (low affinity). As the aminoacyl-tRNA synthetase saturates its primary tRNA target, excess protein binds to the mRNA and inhibits synthesis. It is possible that this structure residing in the first intron or 5′ UTR of the lysyl-tRNA synthetase functions similarly.

#### Metacaspase 1B intronic structure

In our screen of the *N. crassa* genome, we found a structure located within an intron of metacaspase 1B (Fig. 6a). Metacaspases are cysteine-dependent proteases found in protozoa, plants, and fungi. Distantly related to metazoan caspases, they are also involved in programmed cell death and stress pathways [57]. Though caspases and metacaspases have not been reported to have any RNA-binding activity, they are tightly regulated because of their role in apoptosis [58]. A recent study showed post-transcriptional control of executioner caspases in *C. elegans* by RNA-binding proteins [59], and it may be possible that this intronic structure regulates metacaspase 1B upon recognition by an RNA-binding protein.

**Figure 6:**
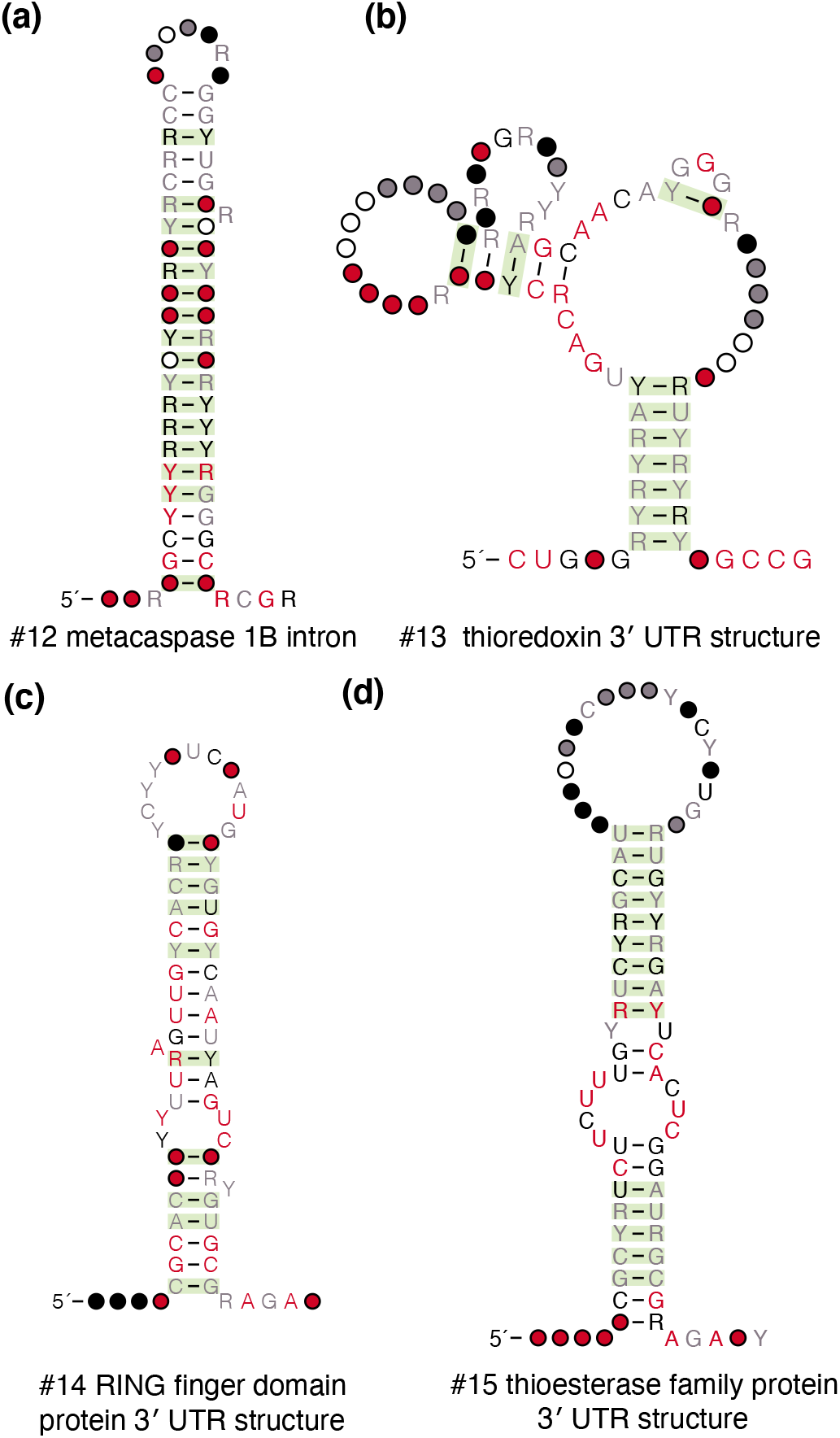
Four structures associated with other mRNAs. Structures were also found in the noncoding regions of these protein-coding genes: **(a)** a structure in an intron of metacaspase 1B, **(b)** a 3′ UTR structure of thioredoxin, **(c)** a 3′ UTR structure in a poorly characterized gene annotated as a RING finger domain protein, and **(d)** a 3′ UTR structure in a poorly characterized gene annotated as a thioesterase family protein.

#### Thioredoxin 3′ UTR structure

In our screens of *N. crassa* and *A. fumigatus,* we also discovered a 3′ UTR structure located in *TXN,* the gene encoding thioredoxin (Fig. 6b). Thioredoxin is an essential protein involved in redox signaling that is found in all living organisms [60]. Interestingly, Yang et al. showed in human colon carcinoma cells that the heterogenous ribonucleoprotein A18 binds to the 3′ UTR of thioredoxin [61]. Although our structure is found in fungi, it is possible that it also interacts with RNA-binding proteins to post-transcriptionally regulate thioredoxin.

#### Two structures in 3′ UTRs of uncharacterized genes

In our screen of *A. fumigatus,* we found two 3′ UTR structures located in poorly characterized genes. One of these is located in a gene annotated by NCBI as encoding a “RING finger domain protein” (Fig. 6c). Although the function of this protein is unclear, it is known that several RING finger domain proteins are involved in ubiquitination [62, 63, 64] and have RNA-binding capabilities [65, 66, 67], suggesting that it may bind to its own RNA. The second structure is located in a gene annotated as a “thioesterase family protein” (Fig. 6d). No thioesterase with an RNA-binding domain has been reported, and a search of the protein sequence against Pfam [68] also did not find any RNA-binding domains in this protein. Therefore, it is possible that other RNA-binding proteins may interact with this structure to regulate protein expression.

#### Two structures in 3′ UTRs of genes encoding metabolic enzymes

Finally, we discovered two 3′ UTR structures in mRNAs encoding metabolic genes. Consistent genomic context across all representatives as well as available RNA-seq data from *S. cerevisiae* and *C. albicans* support this (Figs. 7c-d). In our screen of *S. cerevisiae,* we found the *GLY1* RNA structure, which has 75 representatives within the family Saccharomycetaceae (Fig. 7a). *GLY1* encodes a threonine aldolase that converts the amino acid L-threonine to glycine [69, 70]. *GLY1* is one of the major sources of glycine biosynthesis in *S. cerevisiae* with the other being the *SHM1* /*SHM2* pathway that converts L-serine to glycine; deletion of all three genes is required to render the organism auxotrophic for glycine [71]. Our structure has 16 structurally conserved base pairs supporting all 4 proposed stems which are connected around a multi-stem junction.

**Figure 7:**
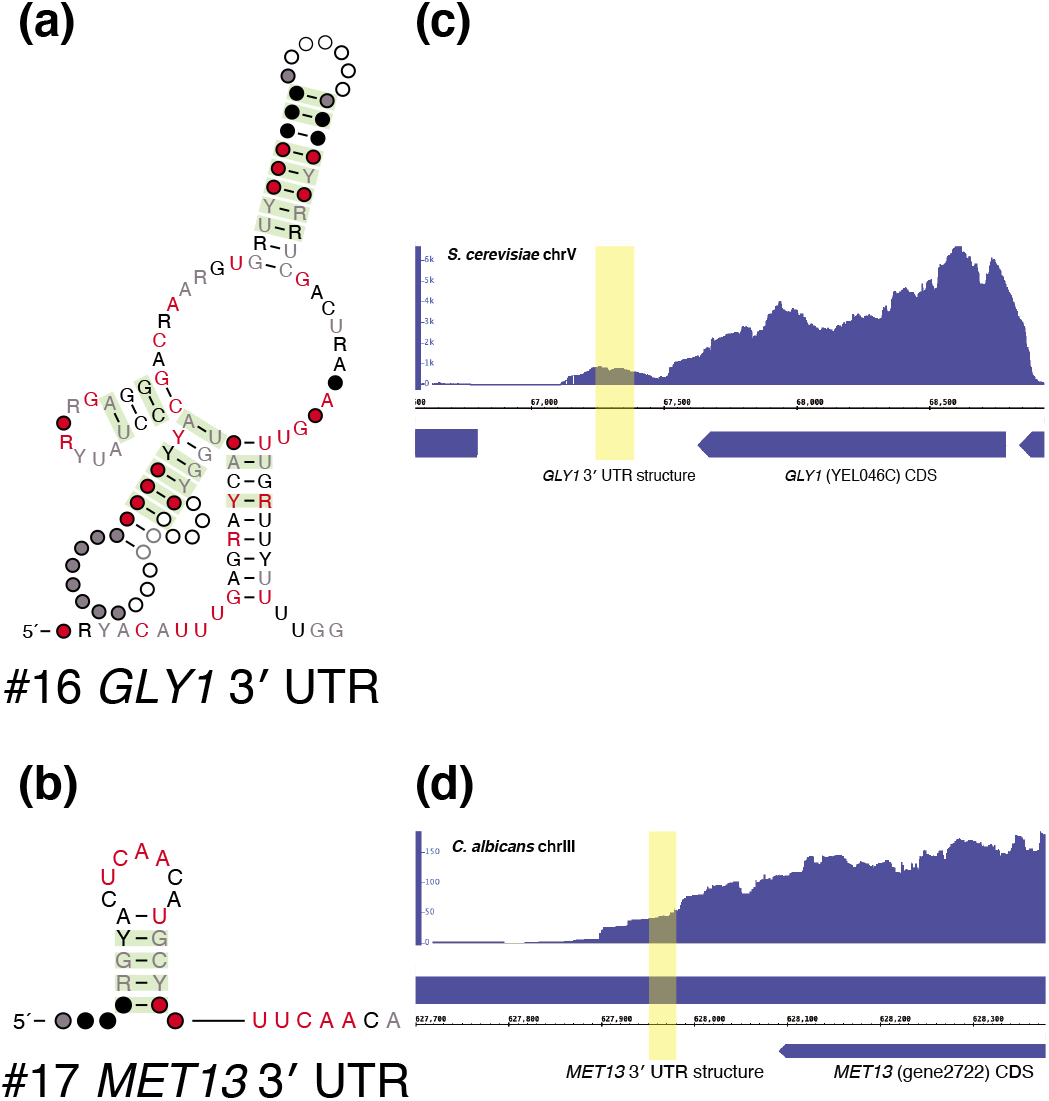
Two eukaryotic 3′ UTR structures. Two structures were found in metabolic genes, which led to us hypothesize that they were eukaryotic riboswitches. These are **(a)** a 3′ UTR structure in *GLY1,* which encodes a threonine aldolase that converts L-threonine to glycine **(b)** a 3′ UTR structure in *MET13,* which encodes methylenetetrahydrofolate reductase, a crucial enzyme in both the folate and methionine cycles. **(c)** RNA-seq data from NCBI Short Read Archive (https://www.ncbi.nlm.nih.gov/sra) accession SRX9766847 was aligned to the *S. cerevisiae* S288C genome using RNA STAR under default settings. Here, the reads aligned to the structure (highlighted in yellow) suggest that it is located in the 3′ UTR of *GLY1.* **(d)** RNA-seq data from SRX6491142 was aligned to the *C. albicans* genome using RNA STAR under default settings. Here, the reads aligned to the structure suggest that it is located in the 3′ UTR of *MET13.* Genome tracks were viewed using IGB [74].

In our screen of *C. albicans,* we discovered a 3′ UTR structure in *MET13,* a gene encoding methylenetetrahydrofolate reductase (Fig. 7b). This enzyme is critical for both the folate cycle, which is involved in purine biosynthesis, and is the rate-limiting step of the methionine cycle, which produces S-adenosyl-L-methionine (SAM), a cofactor that transfers methyl groups to numerous other substrates in enzymatic reactions [72, 73]. Our structure, found in 61 representatives across Debaryomycetaceae/Metschnikowiaceae, is a short hairpin supported by 4 structurally conserved base pairs.

The presence of these structures in the non-coding regions of mRNAs encoding metabolic genes piqued our interest because riboswitches are a well-known class of non-coding cis-regulatory mRNA structures that control metabolic genes by undergoing conformational changes that, without the assistance of any proteins, regulate transcription or translation upon direct and highly specific binding of metabolites. Riboswitches control many bacterial operons [75, 76]. Riboswitches sense a wide array of metabolites, ranging from small ions like fluoride [77] and magnesium [78], small organic compounds including some nucleotides [79] and amino acids [80, 81], to macrocycles like cobalamin (vitamin B_12_) [82].

Despite the abundance of riboswitches sensing diverse substrates in bacteria, only one class of riboswitches has been found in eukaryotes. The TPP riboswitch, located in the 5′ UTR of operons in bacteria [83], is found within a 5′ UTR intron in some fungi and in the 3′ UTR in some plants. In this eukaryotic context, both fungal and plant riboswitches control gene expression by regulating splicing [84, 85].

### 3.6 Experimental characterization of riboswitch candidates

The precedent of 3′ UTR eukaryotic riboswitches led us to experimentally test whether the structures found in the 3′ UTRs of the metabolic enzymes *GLY1* and *MET13* were eukaryotic riboswitches. If so, they would be the first of their kind since the discovery of TPP riboswitches in eukaryotes in 2003 [86]. In bacteria, no threonine aldolase has yet been found under the control of a riboswitch. However, several bacterial genes involved in the glycine cleavage system are controlled by riboswitches responsive to glycine [87]. From a structural standpoint, the *GLY1* 3′ UTR contains a multi-stem junction, a feature common in many known classes of riboswitches [88].

Based on the literature of glycine metabolism in *S. cerevisiae*, we examined whether the *GLY1* 3′ UTR structure was responsive to threonine or glycine, since *GLY1* converts L-threonine to glycine. Additionally, we simultaneously tested several less probable ligands by combining them in a mixture in equal concentrations. These included L-serine and tetrahydrofolate, which are the reactants for *SHM1* and *SHM2*, the other route of glycine biosynthesis, as well as three metabolites that are upstream or downstream of *GLY1* in the glycine metabolic cycle: L-aspartate, adenine, and guanine. *GLY1*’s required cofactor, pyridoxal phosphate (PLP) and TPP, as the only known substrate of a eukaryotic riboswitch, were also included in this mixture.

Though the *MET13* 3′ UTR structure is much less structurally complex, we also decided to test it because some riboswitches including the SAM/SAH riboswitch [14], which binds with high affinity to both SAM and its demethylated form S-adenosyl-L-homocysteine (SAH), are also quite structurally simple. Furthermore, some bacterial homologs of *MET13* are controlled by SAH-sensing riboswitches [89]. Thus, we examined whether the *MET13* 3′ UTR structure was responsive to either SAM or SAH.

To test these hypotheses, we first synthesized a representative *GLY1* 3′ UTR structure from *S. cerevisiae* (164 nts) and a representative *MET13* 3′ UTR structure from *Hyphopichia burtonii* (53 nts) using in vitro transcription. The boundaries of the transcribed sequences matched the boundaries of their conserved structures. We then used these RNAs for in-line probing and isothermal titration calorimetry (ITC) experiments. The in-line probing assay detects RNA-ligand interactions by examining whether an RNA differentially degrades in the presence or absence of a metabolite [37]. Differential degradation suggests that the metabolite induces a change in RNA structure. ITC detects RNA-ligand interactions by measuring the heat absorbed or released upon serial injection of a titrant (a solution containing the metabolite) into a titrate (a solution containing the RNA) [90]. An interaction between the RNA and ligand should yield a thermogram with a sigmoidal curve which represents initial maximal heat of binding followed by eventual saturation once the titrate has been fully bound.

We observed the expected differential degradation in the in-line probing assay and the expected binding thermogram for a positive control, the well-studied *xpt-pbuX* guanine riboswitch from *Bacillus subtilis* (Fig. S3). However, our in-line probing and ITC data for the *GLY1* and *MET13* 3′ UTR structures do not suggest in vitro interactions between either RNA and their corresponding candidate ligands, even when tested at ligand concentrations of 10 mM and 250 *μM* respectively, which is well above the submicromolar dissociation constants of most known riboswitches. No differential degradation was observed in the in-line probing assay and no significant heat of binding was detected between the RNAs and any of the respective candidate ligands for either structure (Figs. S4-S5).

It is still possible that either of these candidates is a riboswitch, and that our in-line probing and ITC experiments are inconclusive because we have not tested the correct ligand. It is also possible that our evolutionary analysis does not show evidence of alternative structures because the expression platform is not included in the alignments. Additional experiments may be needed to further test the riboswitch hypothesis and to propose alternative hypotheses of how these evolutionarily conserved structures are involved in regulating their associated genes.

## 4 DISCUSSION

We have developed a computational method that can take as input any nucleic acid sequence to determine if it has evidence of RNA structural conservation. An iterative nhmmer homology search is used to produce an alignment. This multiple sequence alignment is the input to R-scape, which identifies covariation consistent with conserved RNA structure. While some previous structural RNA gene finding tools also use comparative genomics when predicting RNA structures, R-scape importantly only evaluates conservation of structure using a pure covariation statistic, which quantifies the sole evolutionary pattern unique to structural RNAs: compensatory base pair substitutions. Furthermore, because phylogenetic relationships between sequences may also result in some degree of covariation, R-scape crucially only reports base pairs that covary above this phylogenetic background. In our method, we use the default E-value of 0.05, meaning that we expect fewer than 0.05 apparently false positive base pairs per alignment.

We used R-scape to determine regions that are likely to contain structural non-coding RNAs. To devise appropriate criteria, we first ran our method on several known structural RNAs and used their covariation results to establish a basis for designating an unannotated region as containing a candidate structural ncRNA. We believe that these criteria, namely that the alignment contain at least 3 statistically significant covarying base pairs with at least 2 within the same stem, are quite stringent. Indeed, of over 63,000 intergenic windows screened with the unflanked mode, and over 9,000 introns and over 62,000 UTRs screened with the flanked mode across the five genomes (see Table S1), we identified only 17 novel regions (plus 4 others discovered recently [40, 24, 26]) satisfying our two criteria. Based on the positive control RNAs, we have estimated the expected number of false positives to be 0.003 with 78% (62/80) rate of detection for our set of known structural RNAs in *S. cerevisiae*. It is likely that with laxer criteria we would have identified more candidate structures that could be bona fide structural RNAs, but we selected these criteria to identify only genomic regions with the strongest evidence for structural conservation. We believe these candidates should be priorities for experimental validation.

It is important to note that while these candidates have evolutionary evidence of including conserved RNA structures, their functions can only be determined through experimentation. In all cases, the 5′ and 3′ ends in our alignments are not necessarily the transcript start and end sites, which must be determined experimentally and may differ across organisms. For the structures believed to be associated with mRNAs, the exact mRNA boundaries will be needed to confirm that the structures are located in the UTRs. For the intronic structures, experiments must be done to determine whether they are cis-regulatory elements or standalone genes, like the intronic H/ACA snoRNA we report here. For hypothesized cis-regulatory elements, the protein encoded by the mRNA may present some initial clues as to the possible mechanism of regulation. Here, we used the functions of the GLY1 and MET13 proteins to hypothesize that their mRNA-associated structures were eukaryotic riboswitches. Though the data do not support this, it is interesting to note that they, along with three other structures we identified, are located in the 3′ UTR. Though 3′ UTRs have often been identified as the binding sites of microRNAs and RNA binding proteins, recent studies have revealed unexpected functions of this region [91]. Perhaps future investigation of some of these candidates will expand our understanding of the functional repertoire of 3′ UTRs.

Other methods have used R-scape as a component of their program to identify novel structural ncRNA candidates in viruses [92] and bacteria [93, 94]. The novelty of our method involves the integration of the following approaches: **(a)** Iterative searches and the flanking mode to facilitate primary sequence homology. **(b)** The use of R-scape to identify candidates. Other methods use R-scape either to validate candidates or as one of several approaches. While R-scape extracts only the structural evolutionary signal, other approaches that also measure conservation or consistency with a structure can result in false positives. [95] **(c)** Use of known structural RNAs as positive controls to determine classification criteria based on a desired sensitivity rate (number of positive controls detected) and specificity rate (number of sequences tested multiplied by the highest combined p-value from the detected positive controls). **(d)** Low false positive rate (calculated to be 0.003) in a fungal search.

Our method could be used to study any genome of interest, provided that is well-annotated like the fungal ones we screened here. R-scape’s tunable false positive rate makes it attractive for studying larger eukaryotic genomes, including those of plants and animals.

## 5 Supplemental Data

Supplemental Data are available at a tarball at NAR Online. Supplemental Data include alignments of the 17 structural RNAs, five supplemental figures, three supplemental tables and additional information such as the sequences used for experiments. They also include code to run the flanked and unflanked modes of the method.

## 6 ACKNOWLEDGEMENTS

We thank Michelle Meyer, Alexander Serganov and Ronald Breaker for advice on experimental techniques to characterize riboswitches. We thank Victoria D’Souza and Vladimir Denic for sharing facilities with us to perform the experiments presented in this manuscript. We thank Samuel Carlson, Jerome Edwards and Houqing Yu for numerous experimental tips. We thank Sean Eddy for comments on the manuscript.

## 7 FUNDING

The authors received no specific funding for this work.

## 7.0.1 Conflict of interest statement

None declared.

---

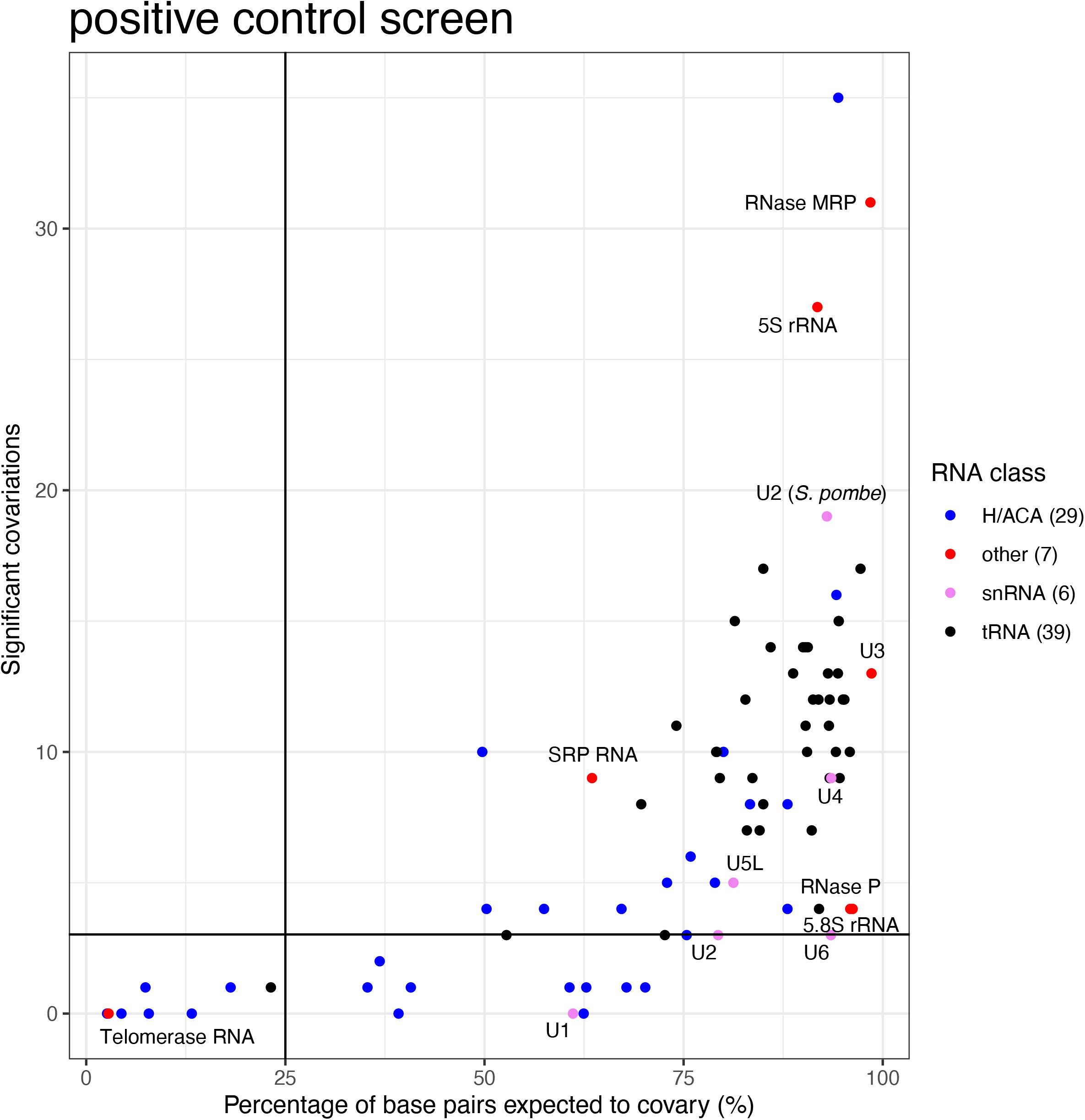

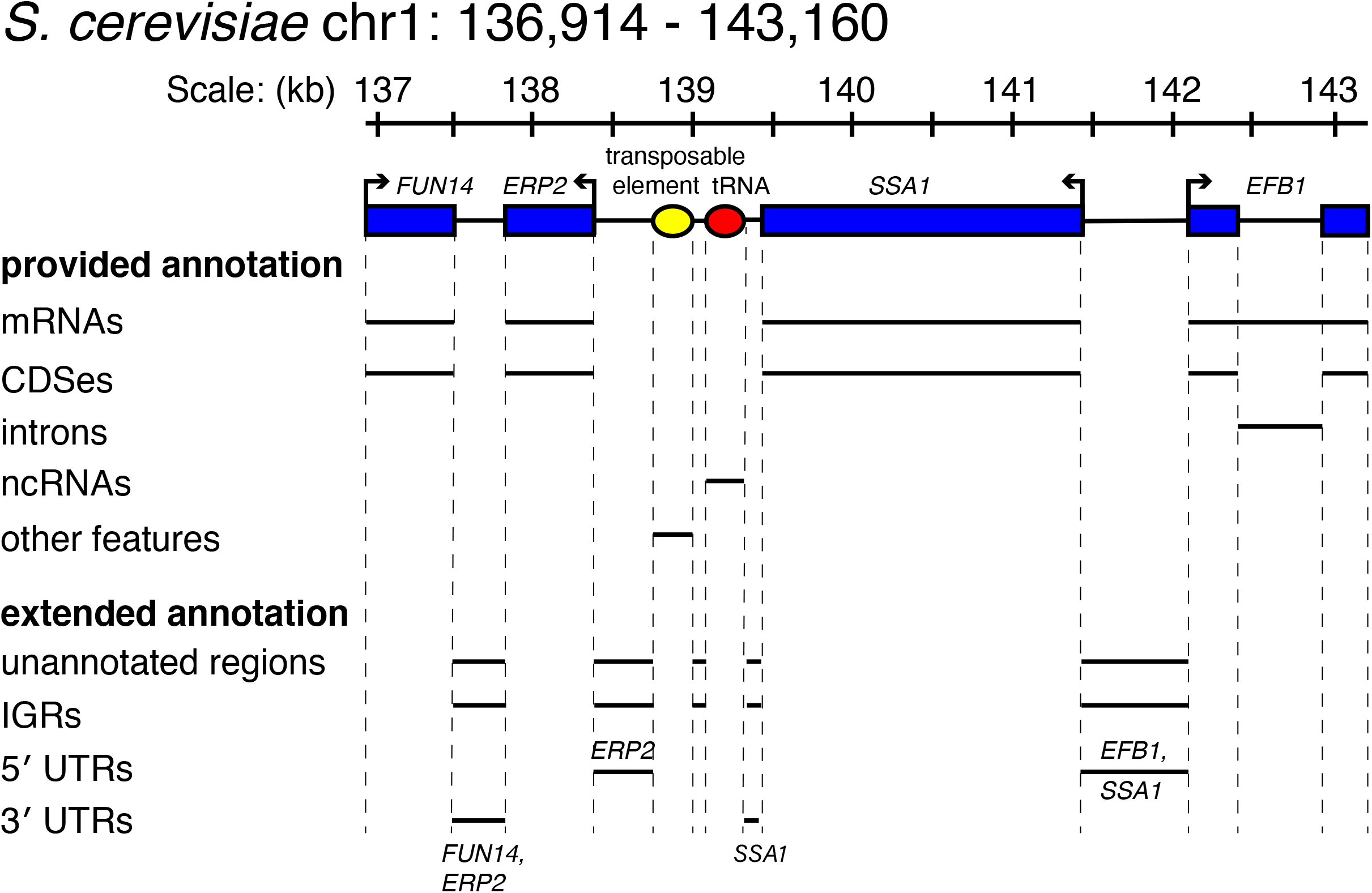

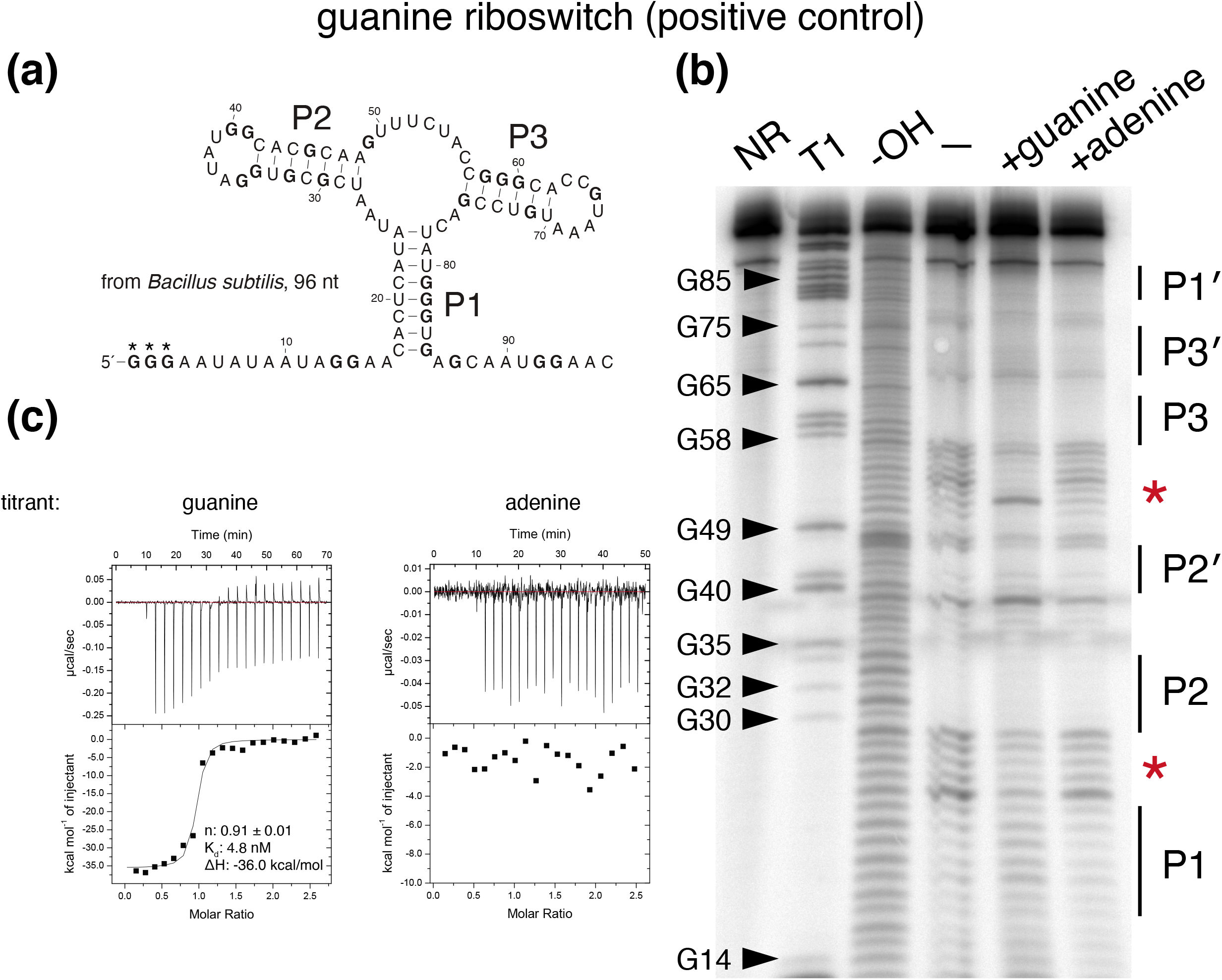

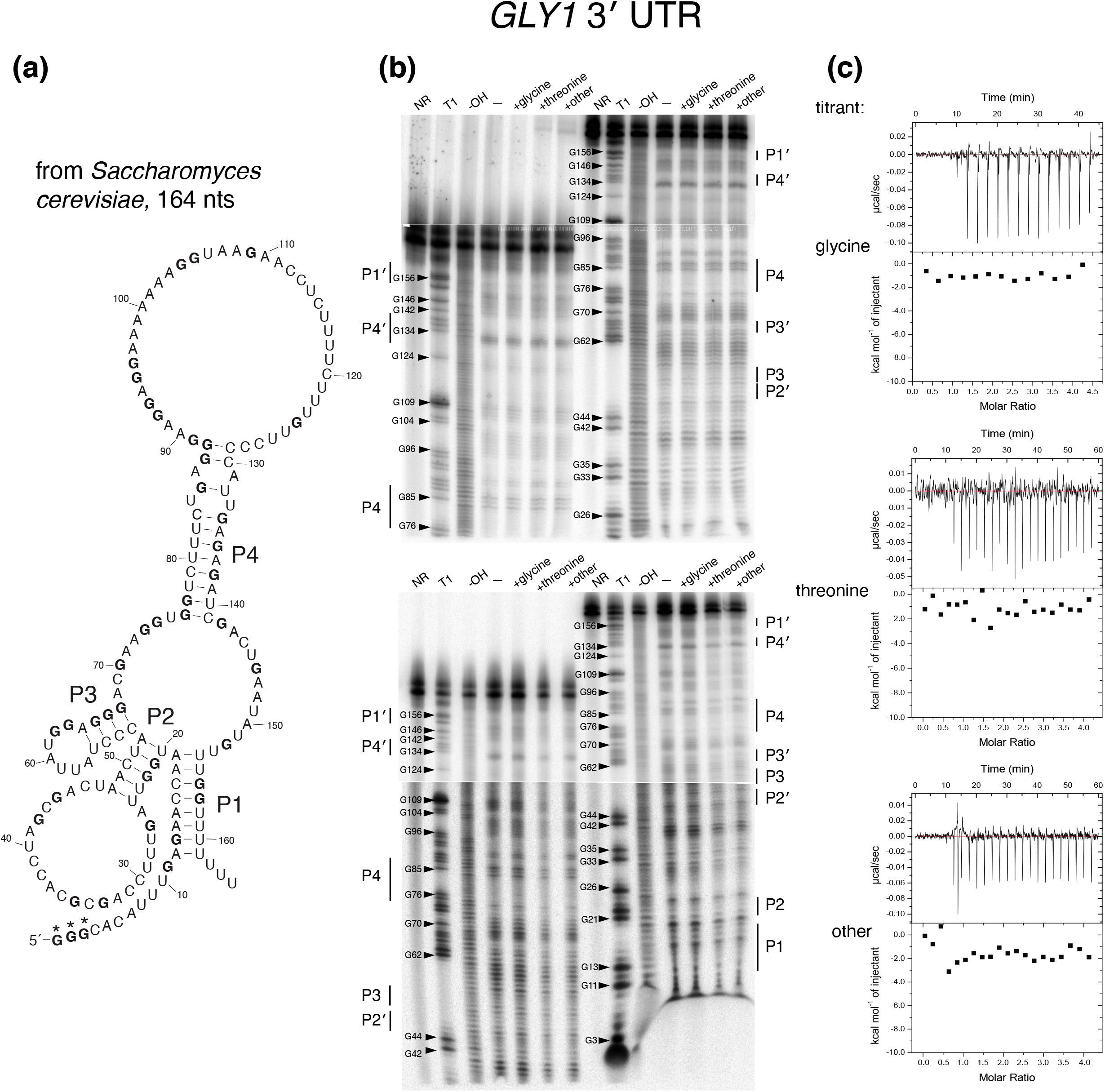

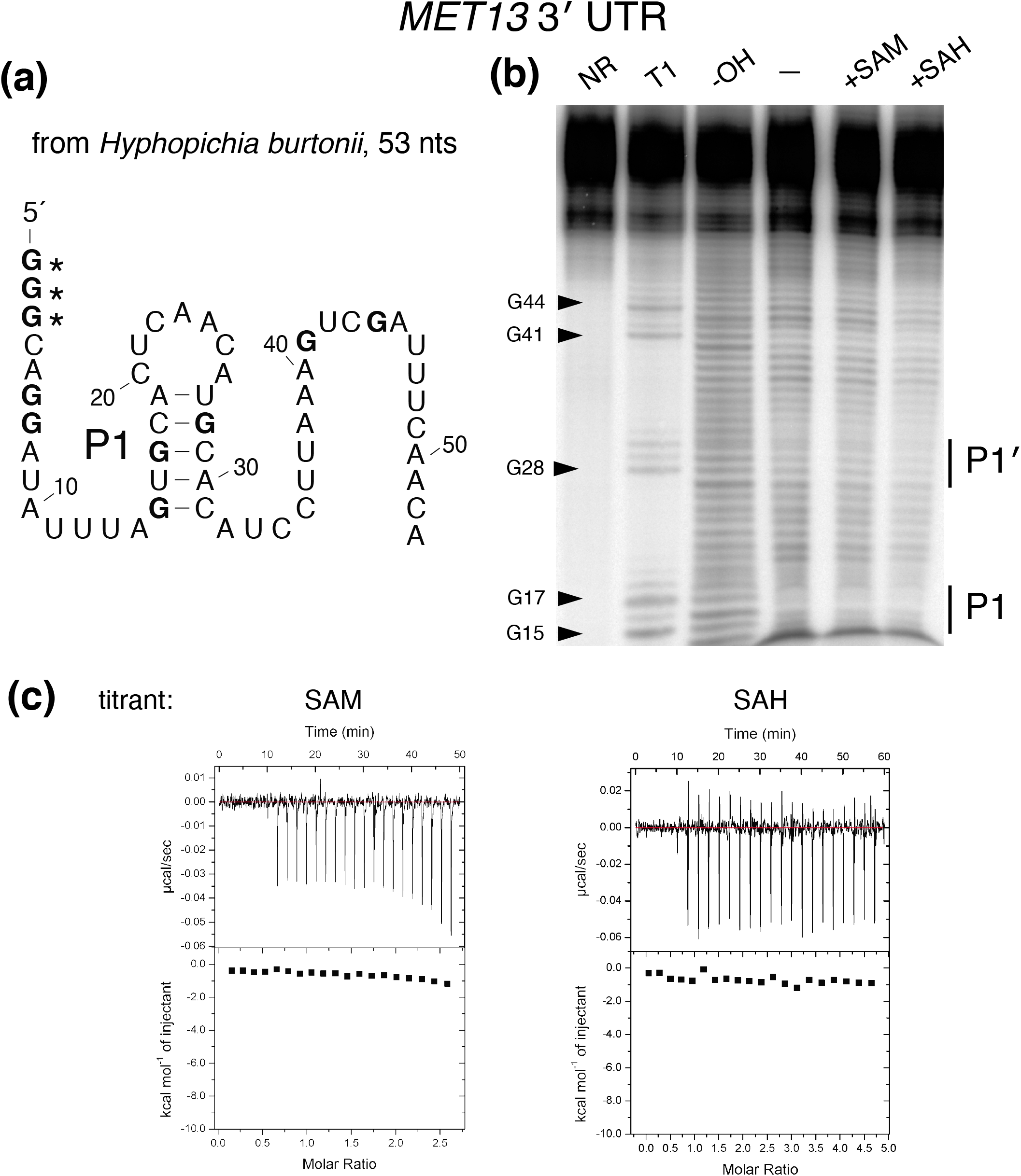

## Notes

http://rivaslab.org

### Competing Interest Statement

The authors have declared no competing interest.

### Summary of Updates

some text edits for clarity.

