## Supplementary material for "Discovery of 17 conserved structural RNAs in fungi": 00README.txt

# Supplemental Data includes

list_of_1371_fungal_genomes.txt       - List of the 1371 genomes in the sequence database

alignments/                           - Folder containing alignments of the 17 structural candidates, in Stockholm format
     nhmmer_alignments/               - Final nhmmer alignments
     infernal_models/		      - Infernal covariance models
     final_alignments/		      - Final structural alignments
     smaller_db_alignments/           - For mRNA-associated candidates 09-17, analysis againgst a smaller db that includes
                                        the mRNA and flanking regions.

methods_scripts/                      - scripts to run the pipeline
        00README
	flanked_mode.sh
	unflanked_mode.sh
	keep_top_genome_hit_only.sh
	subscripts/
	tutorial/
	
supplement/                           - supplemental file with additional methods
    supplement.pdf
