## Supplementary material for "Discovery of 17 conserved structural RNAs in fungi": 00README

Discovery of 17 conserved structural RNAs in fungi
William Gao, Thomas A. Jones, and Elena Rivas

TUTORIAL OF THE FLANKED AND UNFLANKED MODES FOR STRUCTURAL ncRNA DISCOVERY

Step 1: Install software programs

1) Install nhmmer, which is included in the HMMER software package. Download the tar archive from http://hmmer.org/
Per the User Manual, install using the following commands. 
> tar xf hmmer.tar.gz
> cd hmmer-3.3.2
> ./configure
> make
> make install

You will also need to install the Easel commands as part of your installation, using this:
> cd easel; make install

2) Install R-scape. Download the tar archive from http://rivaslab.org/software.html. Per the User Manual, 
install using the following commands:

> tar xf rscape.tar.gz
> cd rscape_v1.5.16
> ./configure
> make
> make install


Step 2: Add paths of software programs to flanked and unflanked mode scripts.

1) The unflanked mode is run using the script unflanked_mode.sh. Insert the path to your nhmmer executable.
2) The flanked mode is run using the script flanked_mode.sh. Insert the path to your nhmmer, esl-alimask, and
esl-alimanip executables.


Step 3: Run examples in the tutorial directory.

To demonstrate usage of the flanked and unflanked mode scripts, a tutorial with working examples is found in the
tutorial/ subdirectory. 

As described in the paper section titled "Method to identify conserved RNA structures" there are two modes of our
structural RNA discovery method. 

1) Unflanked mode:

The unflanked mode is used to screen intergenic regions (IGRs) to discover standalone structural non-coding RNA genes. 
The unflanked mode requires two arguments, in this order. 
>$1	The first argument is a sequence in FASTA format that you wish to run the method on. 
>$2	The second argument is the multi-sequence FASTA file that you wish to screen the query sequence against 
	for homologs (e.g. a genome database).

In the tutorial, a sample IGR region in the Saccharomyces cerevisiae genome is provided. The tutorial uses a small
database of 10 Saccharomyces genomes downloaded from NCBI Genbank in the file tutorial/genome_db/Saccharomyces_genomes.fa. 
Information about the genomes included is available in tutorial/genome_db/genomes_included_in_db.txt.

Using these two arguments, the unflanked mode script can be run as follows:
./unflanked_mode.sh tutorial/IGR_only.fasta tutorial/genome_db/Saccharomyces_genomes.fa

The unflanked mode script produces the following:
>	A directory with the basename of the query sequence, in the same directory as the query sequence. The 
	directory contains the following files:
> 	v1.sto, v2.sto, v3.sto: These are the alignments produced for the first, second, and third iterations of nhmmer.
	Note that the E-values have been set using the -E flag to 1e-10, 1e-10, and 1e-05, as specified in the paper. 
>	v1.hmmout, v2.hmmout, v3.hmmout: These are the tab-delimited files detailing the nhmmer hits. 

2) Flanked mode: 

The flanked mode is used to screen introns and untranslated regions (UTRs) to discover RNA structures associated with mRNAs. 
The flanked mode requires four arguments. 
./flanked_mode.sh <query hmmfile|alignfile|seqfile> <target seqfile (e.g. genome database)> <genome unique string> <noncoding threshold>
>$1	The first argument is a sequence in FASTA format that you wish to run the method on. This sequence should contain 
	non-coding sequence flanked by some amount of protein-coding sequence, since the flanked mode uses the 
	protein-coding sequence to aid homology search in the first nhmmer search. The description field of the 
	FASTA sequence must have its 4th and 5th fields indicate which nucleotides in the sequence correspond to the non-coding region.
	For example, one of the tutorial files has the following descriptor line:
	>IGR_plus_YAR015W_flank chrI 168872-170295 1 503 Saccharomyces cerevisiae S288C
	"IGR_plus_YAR015W_flank" refers to the sequence name, "chrI 168872-170295" gives the genomic location of the 
	full sequence. "1" and "503" indicates that nucleotides 1-503 in this sequence correspond to the non-coding 
	region (i.e. the IGR), while nucleotides 503 to the end (1424) corresponds to the protein-coding flank (i.e.
	the coding sequence of YAR015W).
>$2	The second argument is the multi-sequence FASTA file that you wish to screen the query sequence against for 
	homologs (e.g. a genome database). 
>$3	The third argument is a genome-unique string. The script subscripts/bp_col.sh uses this to identify the columns of 
	the flanked alignment from the first nhmmer search to extract the coordinates corresponding to only the 
	non-coding region. It must be a string that is unique to the genome of the query sequence, and present in the 
	description fields of that genome in the database (For this reason, we recommend that you include an example
	copy of the genome for which you are using as a query. For example, in the tutorial genome_db, we wish 
	to extract the columns of the multi-sequence alignment corresponding to nucleotides 1 to 503 of the S. cerevisiae 
	sequence. One such unique string to do this would be "S288C". 
>$4	The fourth argument is the number of nucleotides that you require the non-coding region to contain after the flanked
	alignment. The threshold we used is 50, meaning that once you extract the non-coding region, only sequences with
	at least 50 nucleotides in this region will be kept to build a HMM. This HMM is then the query for three more 
	iterative homology searches. 

In the tutorial, there are two flanked query sequences provided. They contain the same IGR as in the unflanked mode, flanked by the
upstream and downstream protein-coding sequences. Because the official S. cerevisiae S288C genome annotation annotates
protein-coding genes based only on their CDSes and does not annotate UTRs, the "IGRs" likely contain true intergenic region,
along with the UTRs of the two protein-coding genes. In both alignments here, the entire IGR is treated as if it could potentially
be the UTR of each gene, since it is only 503 nt. For much longer IGRs, it is reasonable to only include intergenic sequences some
distance away from the CDS (e.g. 2000 nt). 

Using these four arguments, the flanked mode script can be run as follows:
./flanked_mode.sh tutorial/YAR014C_plus_IGR.fasta tutorial/genome_db/Saccharomyces_genomes.fa	S288C	50
./flanked_mode.sh tutorial/YAR015W_plus_IGR.fasta tutorial/genome_db/Saccharomyces_genomes.fa	S288C	50

The flanked mode script produces the following:
>       A directory with the basename of the query sequence, in the same directory as the query sequence. 
	The directory contains the following files:
>	flanked.sto, flanked.hmmout: The files produced for the flanked search.
>	noncoding.sto: Only the non-coding region of flanked.sto.
>	trim_noncoding.sto: Removes sequences from noncoding.sto that have fewer than the threshold specified ($4).
>	1_trim.sto, 1_trim.hmmout, 2_trim.sto, 2_trim.hmmout, 3_trim.sto, 3_trim.hmmout: The alignments and tab-delimited result files 
	produced for the first, second, and third iterations of nhmmer on the HMM produced from trim_noncoding.sto. 


Step 4: Optional step (keeping only the top hit per genome)
If you wish to keep only the top hit per genome in any alignment, you can use keep_top_genome_hit_only.sh. This script uses 
esl-alimanip, so edit the path to your executable near the top of the script. This script requires three arguments. 
./keep_top_genome_hit_only.sh <alignment> <outfile name> <genome accession key>
>$1 	The first argument is the alignment for which you want to keep only the top hit per genome.
>$2	The second argument is the name of the outputted alignment. It will be produced in the same directory as the first argument.
>$3 	The third argument is a two-field file, with the first field corresponding to the accessions and the second field corresponding
	to the genome. We recommend that you create this file when concatenating individual genome files into our multiple genome database.
	An example for the tutorial genome database is found here: tutorial/genome_db/genome_accession_key.txt

Once you have finished running the unflanked mode on tutorial/IGR_only.fasta, you should have a directory tutorial/IGR_only
with the final nhmmer alignment called v3.sto. To keep only the top hit per genome, run:
./keep_top_genome_hit_only.sh tutorial/IGR_only/v3.sto rm_v3.sto tutorial/genome_db/genome_accession_key.txt
Once you have finished running the flanked mode on tutorial/YAR014C_plus_IGR.fasta and tutorial/YAR015W_plus_IGR.fasta, you should have
the directories tutorial/YAR014C_plus_IGR/ and tutorial/YAR015W_plus_IGR/ with the final nhmmer alignment called 3_trim_noncoding.sto. To 
keep only the top hit per genome, run:
./keep_top_genome_hit_only.sh tutorial/YAR014C_plus_IGR/3_trim_noncoding.sto rm_v3.sto tutorial/genome_db/genome_accession_key.txt
./keep_top_genome_hit_only.sh tutorial/YAR015W_plus_IGR/3_trim_noncoding.sto rm_v3.sto tutorial/genome_db/genome_accession_key.txt

These scripts will produce a file called "rm_v3.sto" in the same directory as the alignments, with only the top hit per genome included. 


Step 5: Run R-scape on the final alignment
Once you have produced the alignments, you can run R-scape as follows:
R-scape --fold <alignment>

The --fold flag is used because nhmmer alignments do not have a secondary structure prediction. Use R-scape -s --fold if you are running
an alignment with a SS_cons line that contains the secondary structure annotation. Use --outdir if you want to specify a directory for
the R-scape output. For more information on the flags that can be used, run R-scape -h.
