## Supplementary material for "Discovery of 17 conserved structural RNAs in fungi": flanked_mode.sh

```
#!/bin/bash

######################################################################
# RUNS FULL FLANKED MODE, UP TO GENERATION OF FINAL NHMMER ALIGNMENT 
######################################################################

if [ "$#" -ne 4 ]; then
        echo "Illegal number of parameters"
        echo "Must be ./flanked_mode.sh <query hmmfile|alignfile|seqfile> <target seqfile (e.g. genome database)> 
		<genome unique string> <noncoding threshold>"
        exit 1
fi

# software path: edit these accordingly based on your HMMER installation
nhmmer=nhmmer
eslalimask=esl-alimask
eslalimanip=esl-alimanip

# subscript paths
bp_col=subscripts/bp_col.sh
noncoding_extractor=subscripts/noncoding_extractor.sh
noncoding_align=subscripts/noncoding_align.sh

# variables for arguments
query=$1
DB=$2
seq_spec_string=$3
threshold=$4

DIR=$(dirname $query)
filename=$(basename -- "$query")
filename="${filename%.*}"

rm -rf $DIR/$filename
mkdir $DIR/$filename

# first nhmmer search, which should include the CDS flanks
$nhmmer -E 1e-10 -A $DIR/$filename/flanked.sto --tblout $DIR/$filename/flanked.hmmout $query $DB 

# convert columns to nucleotides
$bp_col $DIR/$filename/flanked.sto $seq_spec_string

# extract non-coding region only
$noncoding_extractor $query $eslalimask
rm $DIR/$filename/bp_col.txt 

# align flanking region
$noncoding_align $query $DB $threshold $nhmmer $eslalimanip
```
