## Supplementary material for "Discovery of 17 conserved structural RNAs in fungi": keep_top_genome_hit_only.sh

```
#!/bin/bash

####################################
# KEEPS ONLY THE TOP HIT PER GENOME
####################################

# Check that the correct parameters are given
if [ "$#" -ne 3 ]; then
   	echo "Illegal number of parameters"
	echo "Should be ./keep_top_genome_hit_only.sh <alignment> <outfile name> <genome accession key>"
	exit 1
fi

# software path: edit this accordingly based on your HMMER installation
eslalimanip=esl-alimanip

# variables for arguments
alignment=$1 	# alignment file
outfile_name=$2	# output file name
reference_key=$3  # two-field key file that matches accession numbers to genomes

# directory that all files will be made in
DIR=$(dirname $alignment)
basename=$(basename $alignment)

# temporary files: these are made during the run and will be removed before completion
sequence_map_tmp=$DIR/$basename.map.tmp
sequences_to_remove_tmp=$DIR/$basename.sequences_to_remove.tmp

# names originally in the second column of #=GS
accession_names=($(grep "#=GS" $alignment | sed 's/\// /g' | awk '{print $2}'))
accession_names_with_coord=($(grep "#=GS" $alignment | awk '{print $2}'))

end_index=$((${#accession_names[@]}-1))

# contains 1 to 1 information about the map
if [[ -s $sequence_map_tmp ]]; then
        rm $sequence_map_tmp
fi

for i in $(eval echo "{0..$end_index}")
do
        val=$(awk '$1=="'${accession_names[$i]}'" {print $2}' $reference_key)
        if [[ "$val" == "" ]]; then
                val="NA"
        fi

	echo -n "$val" >> $sequence_map_tmp
	echo -e -n ' \t'  "${accession_names[$i]}" >> $sequence_map_tmp
	echo -e ' \t' "${accession_names_with_coord[$i]}" >> $sequence_map_tmp
done

# determine which genoems have more than one copy in the alignment
duplicates=($(awk '{print $1}' $sequence_map_tmp | sort | uniq -c | awk '$1!="1" {print $2}'))

num_dup=$((${#duplicates[@]}-1))

# print out the number of sequences that need to be removed
echo "Sequences removed: $(($num_dup+1))"

if [[ $num_dup == "-1" ]]; then
	cp $alignment $DIR/$outfile_name
	rm $sequence_map_tmp
	exit 0
fi

if [[ -s $sequences_to_remove_tmp ]]; then
	rm $sequences_to_remove_tmp
fi

# remove the sequences that do not correspond to the top hit
for j in $(eval echo "{0..$num_dup}")
do
	# get everything but the first line (which corresponds to the top hit); the problem here is that the full accession has the coordinates
	partial_accessions=$(grep -w "${duplicates[$j]}" $sequence_map_tmp | tail -n +2 | awk '{print $3}')

	# there may be more than one duplicate
	num_partials=$((${#partial_accessions[@]}-1))
	
	for k in $(eval echo "{0..$num_partials}")
	do
		# get the full accession by looking back at the alignment itself
		grep "${partial_accessions[$k]}" $alignment | awk '$1~/=GS/ {print $2}' >> $sequences_to_remove_tmp
	done
done

# use esl-alimanip to remove sequences
$eslalimanip --seq-r $sequences_to_remove_tmp $alignment > $DIR/$outfile_name

# remove files
rm $sequence_map_tmp
rm $sequences_to_remove_tmp
```
