## Supplementary material for "Discovery of 17 conserved structural RNAs in fungi": bp_col.sh

```
#!/bin/bash

if [ "$#" -ne 2 ]; then
   	echo "Illegal number of parameters"
	echo "Should be ./bp_col.sh <alignment> <genome specific string>"
	exit 1
fi


# names of files, provided based on first two arguments
alignment=$1
seq_spec_string=$2

# directory of alignment file that you provide, the output will go into there
DIR=$(dirname $alignment)

# Read the alignment file to get the top sequence that contains reference genome 
seq_name=$(grep "$seq_spec_string" $alignment | head -1 | awk '{print $2}')

# Extract only the sequence (gaps included) of that sequence from the MSA
grep $seq_name $alignment | awk '$2!~/\// {print $2}' | tr -d '\n' > $DIR/bar.txt

# Now create a counter that loops through and increments as it reads each character.
while read -n1 c; do
	counter=$((counter+1))
	c=$c
	if [[ $c = *"U"* || $c = *"A"* || $c = *"T"* || $c = *"G"* || $c = *"C"* ]]; then
		
		if [[ $c = *"U"* ]]; then
			c="T"
		fi

		bp[i]=$c
		array[$i]=$counter
		i=$((i+1))
	fi
done < $DIR/bar.txt

# header
        echo -n "bp" > $DIR/bp_col.txt
        echo -n -e ' \t ' >> $DIR/bp_col.txt
        echo "Col" >> $DIR/bp_col.txt

# echo each element of the array
k=1
for each in "${array[@]}"
do
	echo -n $k >> $DIR/bp_col.txt
	echo -n -e ' \t ' >> $DIR/bp_col.txt
	echo "$each" >> $DIR/bp_col.txt
	k=$((k+1))
done

rm $DIR/bar.txt
```
