## Supplementary material for "Discovery of 17 conserved structural RNAs in fungi": noncoding_align.sh

```
#!/bin/bash

#####################################################################
## EXTRACTS AT A CERTAIN THRESHOLD (NUMBER OF NUCLEOTIDES) WITHIN THE
## INTRON/UTR, BY USING ESL-ALIMANIP (TRIM_NONCODING.STO)
## RUN THREE ITERATIONS WITH THAT ALIGNMENT (3_TRIM_NONCODING.STO)
####################################################################

if [ "$#" -ne 5 ]; then
        echo "Illegal number of parameters"
        echo "Must be ./noncoding_align.sh <.fasta file> <DB> <noncoding threshold> <nhmmer path> <esl-alimanip path>"
        exit 1
fi

query=$1
DB=$2
threshold=$3
nhmmer=$4
eslalimanip=$5

DIR=$(dirname $query)
filename=$(basename -- "$query")
filename="${filename%.*}"

# Trim alignment, include only sequences that have at least N nt in the non-coding region, specified by threshold
$eslalimanip --lmin $threshold $DIR/$filename/noncoding.sto > $DIR/$filename/trim_noncoding.sto

# Iterative homology search, using only the non-coding region
$nhmmer -E 1e-10 -A $DIR/$filename/1_trim_noncoding.sto --tblout $DIR/$filename/1_trim.hmmout $DIR/$filename/trim_noncoding.sto $DB 
$nhmmer -E 1e-10 -A $DIR/$filename/2_trim_noncoding.sto --tblout $DIR/$filename/2_trim.hmmout $DIR/$filename/1_trim_noncoding.sto $DB 
$nhmmer -E 1e-05 -A $DIR/$filename/3_trim_noncoding.sto --tblout $DIR/$filename/3_trim.hmmout $DIR/$filename/2_trim_noncoding.sto $DB
```
