## Supplementary material for "Discovery of 17 conserved structural RNAs in fungi": noncoding_extractor.sh

```
#!/bin/bash

##############################################################################################
## EXTRACTS THE NONCODING (INTRON/UNTRANSLATED REGION) ALIGNMENT FROM THE WHOLE GENE ALIGNMENT
##############################################################################################

if [ "$#" -ne 2 ]; then
        echo "Illegal number of parameters"
        echo "Must be ./noncoding_extractor.sh <query FASTA file> <esl-alimask path>"
        exit 1
fi

query=$1
eslalimask=$2

DIR=$(dirname $query)
filename=$(basename -- "$query")
filename="${filename%.*}"

s1=$(awk '$1~/>/ {print $4}' $query)
s2=$(awk '$1~/>/ {print $5}' $query)

first_site=$(awk '$1=="'$s1'" {print $2}' $DIR/$filename/bp_col.txt)
second_site=$(awk '$1=="'$s2'" {print $2}' $DIR/$filename/bp_col.txt)

$eslalimask -t $DIR/$filename/flanked.sto $first_site..$second_site > $DIR/$filename/noncoding.sto
```
