## Supplementary material for "Discovery of 17 conserved structural RNAs in fungi": unflanked_mode.sh

```
#!/bin/bash

######################################################################
# RUNS FULL UNFLANKED MODE, UP TO GENERATION OF FINAL NHMMER ALIGNMENT 
######################################################################

if [ "$#" -ne 2 ]; then
        echo "Illegal number of parameters"
        echo "Must be ./unflanked_mode.sh <query hmmfile|alignfile|seqfile> <target seqfile (e.g. genome database)>"
        exit 1
fi

# software path: edit this accordingly based on your HMMER installation
nhmmer=nhmmer

# variables for arugments
query=$1
DB=$2

# create new directory to store files from unflanked mode
# the directory name is the query sequence file name 
DIR=$(dirname $query)
filename=$(basename -- "$query")
filename="${filename%.*}"
rm -rf $DIR/$filename
mkdir $DIR/$filename

# first iteration, E-value: 1e-10
$nhmmer -E 1e-10 -A $DIR/$filename/v1.sto --tblout $DIR/$filename/v1.hmmout $query $DB

# second iteration, E-value: 1e-10
$nhmmer -E 1e-10 -A $DIR/$filename/v2.sto --tblout $DIR/$filename/v2.hmmout $DIR/$filename/v1.sto $DB 

# third iteration, E-value: 1e-05
$nhmmer -E 1e-05 -A $DIR/$filename/v3.sto --tblout $DIR/$filename/v3.hmmout $DIR/$filename/v2.sto $DB
```
