## Supplementary material for "Discovery of 17 conserved structural RNAs in fungi": supplement.pdf

Table S3. Structures that satisfied the selection criteria but had a significant hit to any Rfam families were removed. The list of 106 removed hits includes: 35 rRNAs, 32 tRNAs, 25 snoRNAs, and 15 other RNA genes.

#### Supplemental Figures

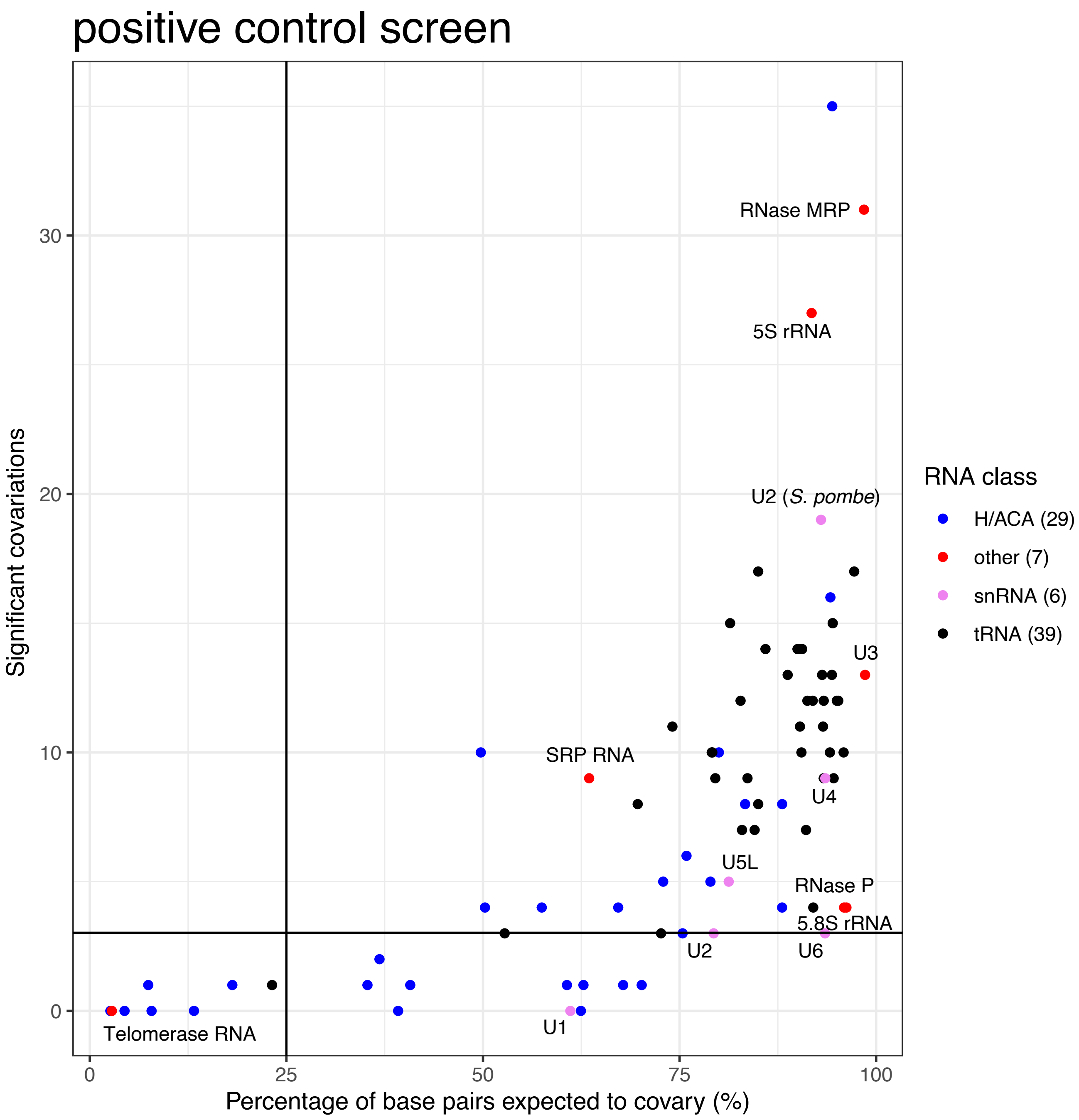


Figure S1: **Results of optimized method in positive controls**. When optimizing our method for discovering structural non-coding RNAs, we tested how two parameters, the number of iterative homology searches and E-value threshold, affected the number of statistically significant covariations (inferred as conserved base pairs) detected in alignments of these known functional ncRNAs. This set of positive controls included 80 structural ncRNA genes from the *S. cerevisiae* genome, including all 29 H/ACA box small nucleolar RNAs (snoRNA), 5S and 5.8S ribosomal RNAs (rRNA), 39 transfer RNAs (tRNA) with one per anticodon, 5 spliceosomal RNAs (snRNA), and 5 other RNAs (RNase MRP, RNase P, SRP RNA, Telomerase RNA, and U3). These sequences were flanked with upstream and downstream intergenic sequences, to simulate a screen for standalone ncRNA genes using the unflanked mode. With the optimized unflanked mode (see Main Fig. 2), 62 out of 80 (78%) ncRNAs have 3 or more conserved base pairs, and only 10 (13%) have fewer than 3 conserved base pairs when more than 25% of base pairs are expected to covary (measure of covariation power). 8 (10%) of these ncRNAs have less than a 25% of base pairs expected to covary, and in all cases, these ncRNAs have low covariation power because homology is only detected in the *Saccharomyces sensu stricto*. In some cases, fewer conserved base pairs are found than expected because nhmmer does not produce alignments that account for secondary structure or because of peculiarities in the query sequence. For example, the U2 sequence from *S. cerevisiae* is unusually large compared to homologs from other fungi (1). Using a sequence more representative of fungal U2s such as the one from *S. pombe* resulted in detection of many more conserved base pairs. All ncRNAs except H/ACA box snoRNAs and tRNAs are labeled on the plot.

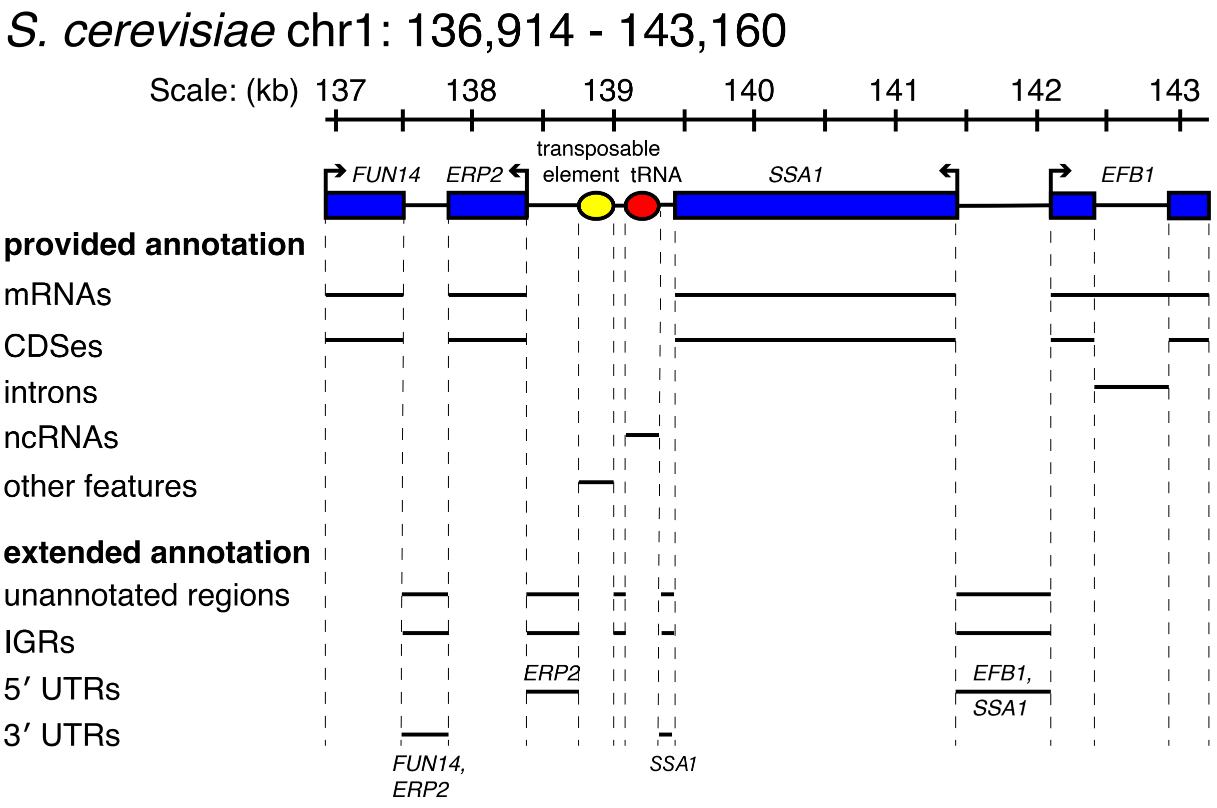


Figure S2: **Example of a genomic region screened**. Here, a 6 kb region of the *S. cerevisiae* genome is shown with annotations from the most recent (01/13/2015) annotation from the *Saccharomyces* Genome Database. The annotations give the coordinates of protein-coding genes, non-coding genes, and other features like transposable elements. The coordinates of introns are also provided. Based on this annotation, we defined intergenic regions (IGRs) as any unannotated regions. If intronic coordinates are explicitly provided (as for *S. cerevisiae* and *S. pombe*), we use those coordinates. In the annotations of the other three query genomes, the intron coordinates are not explicitly defined, so we define them as the intervening sequence between two annotated exons of the same protein-coding gene. If the coordinates of untranslated regions (UTRs) are provided (only for *S. pombe*), then we use those coordinates. In the four other genomes, the IGR coordinates were treated as the UTR coordinates, with the direction of transcription of the adjacent gene used to designate the UTR as either 5′ or 3′.

### Experiments with positive control guanine riboswitch

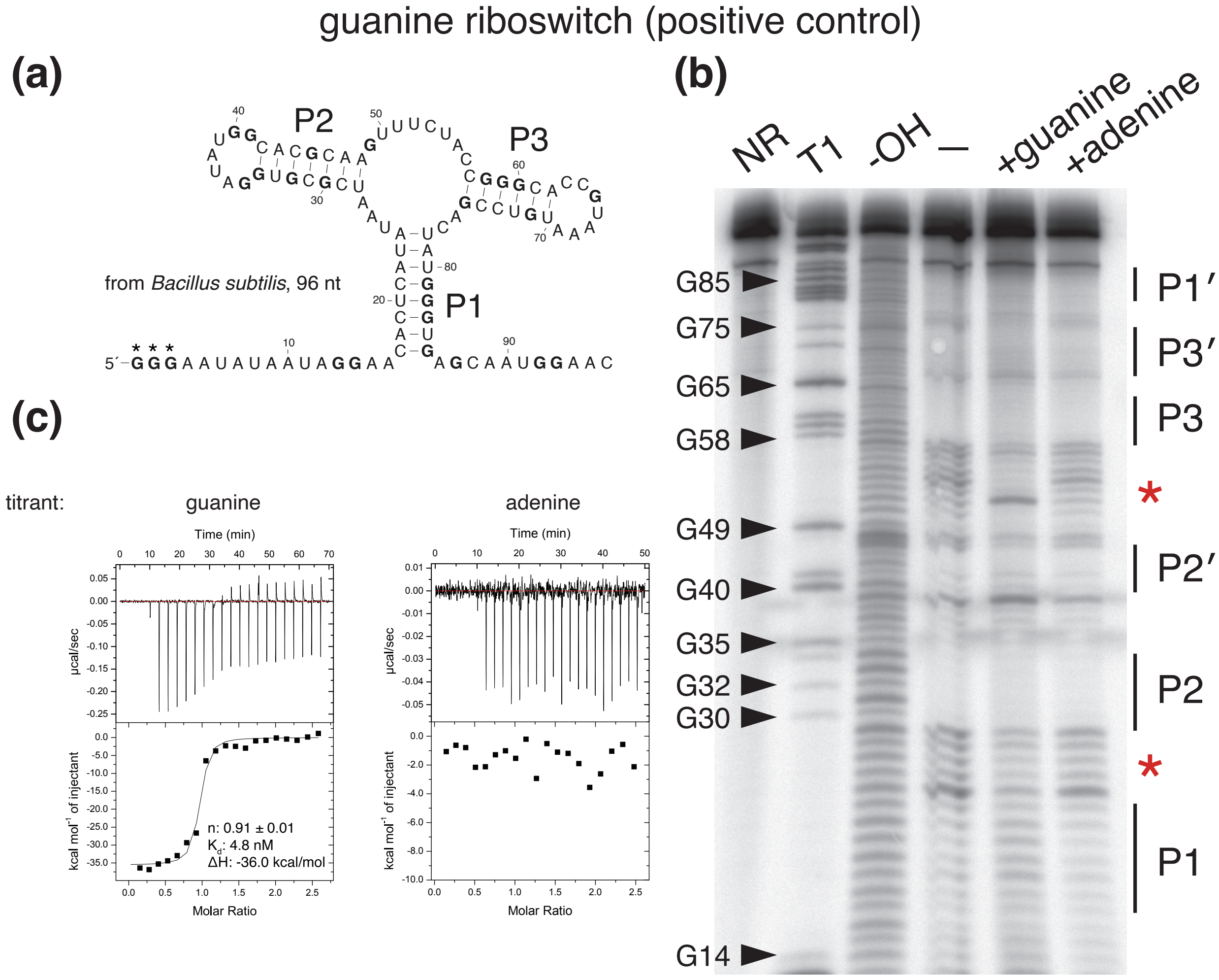


Figure S3: **Experiments with guanine riboswitch**. **(a)** The sequence and structure of the *B. subtilis xpt-pbuX* guanine riboswitch. Guanine residues are bolded. The first three guanine residues (in asterisks) were added to the beginning of the sequence to increase efficiency of in vitro transcription. **(b)** In-line probing of guanine riboswitch: red asterisks indicate the regions of differential degradation between the lane incubated with 10 $\mu$M guanine (+guanine) and those incubated with no ligand (-) or 10 $\mu$M adenine (+adenine). The no reaction (NR) lane corresponds to undigested, precursor RNA, the T1 lane corresponds to the RNA digested with RNase T1 which cleaves after guanine residues, and -OH lane corresponds to the RNA under partial alkaline digestion, which produces a single nucleotide ladder. Typically, structured regions are less susceptible to degradation, and our gel is concordant with this: the base pairing regions labeled "P1" to "P3" are less frequently the site of degradation than unpaired regions. The regions of differential degradation are also consistent with gels from the literature (2). **(c)** Representative isothermal titration calorimetry results for the guanine riboswitch. The top window in each plot shows the raw ITC data and the bottom window shows $\Delta$H. 1.5 $\mu$M RNA was titrated against 15 $\mu$M guanine (left) or adenine (right). Only titration with guanine results in the expected sigmoidal binding isotherm and has derived thermodynamic parameters consistent with the literature (3).

### Experiments with *GLY1* 3′ UTR and *MET13* 3′ UTR

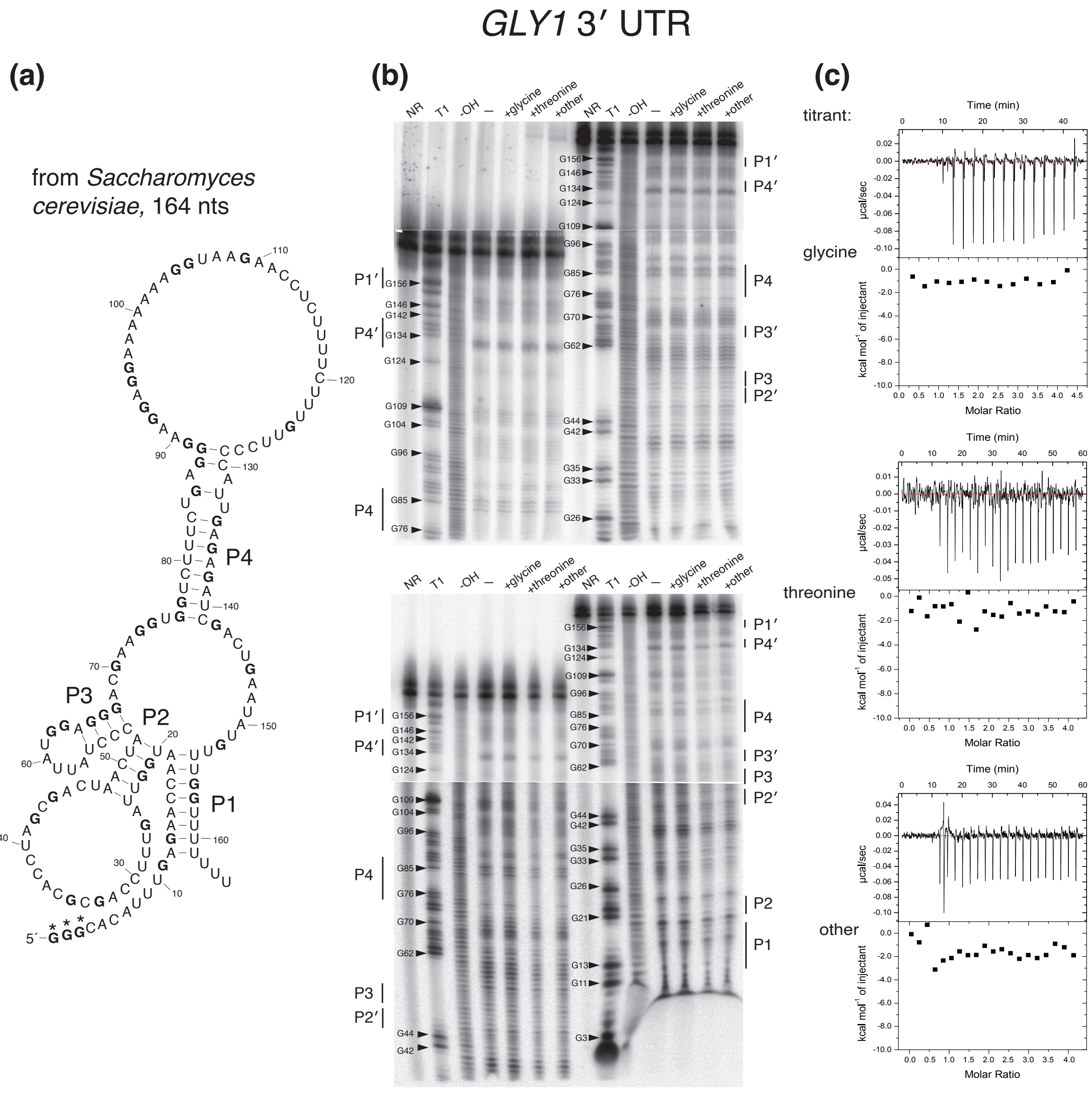


Figure S4: **Experiments with the *GLY1* 3′ UTR structure**. **(a)** The sequence and proposed structure of the *GLY1* 3′ UTR structure from *S. cerevisiae*. Guanine residues are bolded. The first three guanine residues (in asterisks) were added to the beginning of the sequence to increase efficiency of in vitro transcription. **(b)** Two gels from the in-line probing assay are shown. In each case, half of the sample was run initially at 50 W on a sequencing gel. After three hours, the second half was loaded. Therefore, the first half was run for 6 hours total to increase resolution of the 3′ end of the RNA. There is no differential degradation between the RNA incubated with no ligand (-) compared to those incubated with 10 mM glycine, 10 mM L-threonine, or a mixture of several other compounds (L-serine, tetrahydrofolate, L-aspartate, guanine, adenine, PLP, and TPP) all at 10 mM. The no reaction (NR) lane corresponds to undigested, precursor RNA, the T1 lane corresponds to the RNA digested with RNase T1 which cleaves after guanine residues, and the -OH lane corresponds to the RNA under partial alkaline digestion, which produces a single nucleotide ladder. Typically, structured regions are less susceptible to degradation, and our gel is concordant with this: the base pairing regions labeled “P1” to “P4” are less frequently the site of degradation than unpaired regions. **(c)** Representative isothermal titration calorimetry results. The top window in each plot shows the raw ITC data and the bottom window shows $\Delta$H. 5 $\mu$M RNA was titrated against 100 $\mu$M of either glycine, threonine, or the mixture of other ligands. None resulted in the sigmoidal binding isotherm expected if a ligand were binding to the RNA.

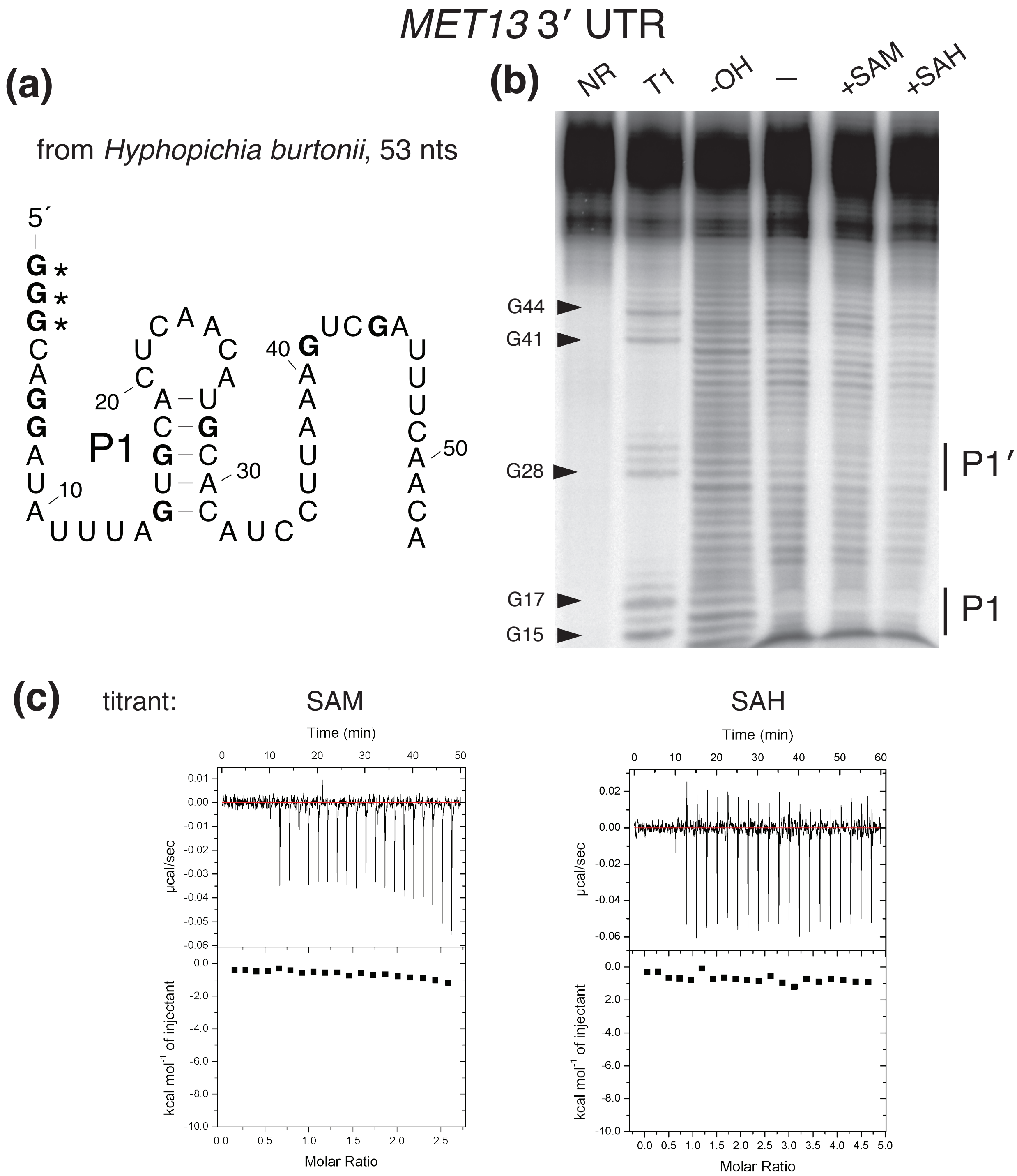


Figure S5: **Experiments with the *MET13* 3′ UTR structure**. **(a)** The sequence and proposed structure of the *MET13* 3′ UTR structure from *H. burtonii*. Guanine residues are bolded. The first three guanine residues (in asterisks) were added to the beginning of the sequence to increase efficiency of in vitro transcription. **(b)** Results from the in-line probing assay are shown. There is no differential degradation between the RNA incubated with no ligand (-) compared to those incubated with 250 $\mu$M SAM or SAH. The no reaction (NR) lane corresponds to undigested, precursor RNA, the T1 lane corresponds to the RNA digested with RNase T1 which cleaves after guanine residues, and the -OH lane corresponds to the RNA under partial alkaline digestion, which produces a single nucleotide ladder. **(c)** Representative isothermal titration calorimetry results. The top window in each plot shows the raw ITC data and the bottom window shows $\Delta$H. On the left, 12 $\mu$M RNA was titrated against 150 $\mu$M SAM and on the right 8 $\mu$M RNA was titrated against 180 $\mu$M SAH. Neither resulted in the sigmoidal binding isotherm expected if a ligand were binding to the RNA.
