## Supplementary material for "Discovery of 17 conserved structural RNAs in fungi": supplement.tex

\documentclass[12pt]{report}
\usepackage[utf8]{inputenc}
\usepackage{graphicx}
\usepackage{gensymb}
\usepackage{pdflscape}
\usepackage{rotating}
\usepackage{longtable}

\usepackage{caption}
\captionsetup{font=footnotesize}

\usepackage[colorlinks = true,
            linkcolor = blue,
            urlcolor  = blue,
            citecolor = blue,
            anchorcolor = blue]{hyperref}

% PACKAGES %
\usepackage[legalpaper, portrait, margin=1in]{geometry}

% headers and footers

% this package allows for including figures in LaTEX 
\usepackage{tgtermes}
\usepackage[font={footnotesize}]{caption}
\usepackage{mathtools} 
\usepackage{amsmath}  
\usepackage{amssymb} 
% do not box figures
\usepackage{float}
\floatstyle{plain}
%\floatstyle{boxed} 
\restylefloat{figure}
% package of subfigures
\usepackage{subcaption}
% customize spacing
\usepackage{setspace}
\linespread{1.5}
\setlength\parindent{24pt}
% remove number of sections and subsections
\setcounter{secnumdepth}{0}

% packages for tables
\usepackage{array}
\usepackage{makecell}

\newcommand{\lb}[1]{\raisebox{-1.0ex}[0pt]{#1}}

\begin{document}

\title{\textbf{Discovery of 17 conserved structural RNAs in fungi}\\ Supplemental Materials}	
\author{William Gao, Thomas A. Jones, and Elena Rivas}

\maketitle


\section{Supplemental Tables}

\begin{table}[h]
  Intergenic regions (unflanked mode)
  
  \bigbreak	
  
  \begin{tabular}{ | c | c | c | c | c | c | }
    \hline
    \thead{Species}                             &
    \thead{Nuclear \\  genome size \\  (Mb)   } &
    \thead{\# IGRs}                             &
    \thead{\# windows \\ under \\ 100 nt}       &
    \thead{\# windows w/o \\ final  \\ nhmmer}  &
    \thead{\# windows w/ \\ final \\  nhmmer}  \\
    \hline
    \textit{S. cerevisiae} & 12.1 &  6699 & 589 &   271 &  7031 \\
    \hline
    \textit{C. albicans}   & 14.3 &  6263 & 639 &  3069 &  7180 \\
    \hline
    \textit{N. crassa}     & 40.5 &  9173 & 555 & 10714 & 22664 \\
    \hline
    \textit{A. fumigatus}  & 29.4 & 11709 & 600 &  2977 & 22484 \\
    \hline
    \textit{S. pombe}      & 12.6 &  4401 & 975 &  178 &   4320 \\
    \hline
  \end{tabular}
  
  \bigbreak
  \bigbreak
  
  Untranslated regions / introns (flanked mode)
  
  \bigbreak
  
  \begin{tabular}{ | c | c | c | c | c | c | c | c  | c | }
    \hline
    \thead{Species}                            &
    \thead{\# protein- \\ coding \\ genes}     &
    \thead{\# introns \\ under \\ 100 nt}      &
    \thead{\# introns w/o  \\ final \\ nhmmer} &
    \thead{\# introns w/ \\ final  \\ nhmmer}  &
    \thead{\# UTRs \\ under \\ 100 nt}         &
    \thead{\# UTRs w/o \\ final \\ nhmmer}     &
    \thead{\# UTRs w/ \\ final \\ nhmmer}  \\
    \hline
    \textit{S. cerevisiae} &  6572 &   110 &  9 &  190 &  870 &  39 & 10748 \\
    \hline
    \textit{C. albicans}   &  6245 &   237 &  0 &  165 & 1227 & 356 & 10612 \\
    \hline
    \textit{N. crassa}     & 10381 & 13908 & 55 & 6044 & 1069 & 517 & 16445 \\
    \hline
    \textit{A. fumigatus}  &  9859 & 16208 &  8 & 2413 &  723 & 797 & 17496 \\
    \hline
    \textit{S. pombe}      &  7097 &   634 &  0 &  236 & 1332 & 158 &  6569 \\
    \hline
  \end{tabular}
  \caption*{Table S1. \textbf{Details of fungal screen.} For each of the five query genomes, we report the number of intergenic regions subject to the unflanked mode, and the number of UTRs and intronic sequences subject to the flanking mode. Query sequences shorter than 100 nt were removed from the analysis, because an exact match of 100 nt was empirically determined for our 1371 fungal genome database to have an E-value close to the E-value threshold of $10^{-10}$ for the first iteration. For the remaining sequences, we report the number with alignments obtained after the 3-fold iterative homology search performed using nhmmer. Almost all of the sequences that do not have a final nhmmer alignment are from low complexity or highly repetitive regions that were not provided in the genome annotations. UTRs and intergenic regions were only defined for regions that are not overlapping with any other annotations. \textit{S. pombe} is an genome where a significant portion of annotations are overlapping, resulting in fewer UTRs and intergenic regions than expected for the number of annotated protein-coding genes. Refer to Fig. S2 for an example of a genomic region and how annotations were used to identify IGRs, introns, and UTRs.}
\end{table}

\pagebreak
\begin{landscape}
  \begin{table}[H]
  \begin{center}
    \small
    \begin{tabular}{|l|l|l|r|l|l|}
      \hline
      \multicolumn{1}{|c|}{\textbf{Structural RNA}} & \multicolumn{1}{|c|}{\textbf{Model Organism(s)}} & \multicolumn{1}{|c|}{\textbf{Coordinates}} & \multicolumn{1}{|c|}{\textbf{Size (nts)}}& \multicolumn{1}{|c|}{\textbf{Closest gene}}\\
      \hline
      \thead[l]{01 H/ACA snoRNA 1}                               & \thead[l]{\textit{C. albicans}}                      & \thead[l]{chr2:424209-424497}                         & \thead[r]{289}      & \thead[l]{SAM37: 422849-423919}	\\
      \hline
      \thead[l]{02 H/ACA snoRNA 2}                               & \thead[l]{\textit{C. albicans}}                      & \thead[l]{chrR:2099614-2100061}                       & \thead[r]{448}      & \thead[l]{FGR46: 2100560-2100198} \\
      \hline
      \thead[l]{03 H/ACA snoRNA 3}                               & \thead[l]{\textit{A. fumigatus}}                     & \thead[l]{chr7:1017260-1017427}                       & \thead[r]{168}      & \thead[l]{AFUA\_7G04460: 1015962-1017200} \\
      \hline	
      \thead[l]{04 H/ACA snoRNA intronic}                        & \thead[l]{\textit{A. fumigatus}}                     & \thead[l]{chr2:3571437-3571242}                       & \thead[r]{196}      & \thead[l]{AFUA\_2G13750: 3571860-3569481} \\
      \hline
      \thead[l]{05 conserved intergenic structure 1}             & \thead[l]{\textit{A. fumigatus}\\\textit{N. crassa}} & \thead[l]{chr3:2027363-2027443\\chr1:8347842-8347751} & \thead[r]{81\\92}   & \thead[l]{AFUA\_3G07970: 2028224-2026400\\NCU03041: 8346947-8349780}  \\
      \hline
      \thead[l]{06 conserved intergenic structure 2}             & \thead[l]{\textit{A. fumigatus}}                     & \thead[l]{chr1:4003534-4003702}                       & \thead[r]{169}      & \thead[l]{AFUA\_1G14940: 4004644-4007079} \\
      \hline
      \thead[l]{07 conserved intergenic structure 3}             & \thead[l]{\textit{A. fumigatus}\\\textit{N. crassa}} & \thead[l]{chr2:899227-899360\\chr2:2737780-2737909}   & \thead[r]{134\\130} & \thead[l]{AFUA\_2G03420: 899213-901323\\NCU00133: 2737574-2740109} \\
      \hline
      \thead[l]{08 conserved intergenic structure 4}             & \thead[l]{\textit{A. fumigatus}\\\textit{N. crassa}} & \thead[l]{chr5:3359121-3359194\\chr5:2294613-2294542} & \thead[r]{74\\72}   & \thead[l]{AFUA\_5G12830: 3360387-3359565\\NCU03749: 2292401-2294704} \\ 
      \hline
      \thead[l]{09 \textit{RPL12} 5$^\prime$ UTR}                 & \thead[l]{\textit{A. fumigatus}}                     & \thead[l]{chr1:988401-988346}                         & \thead[r]{56}       & \thead[l]{AFUA\_1G03390: 988333-987678} \\
      \hline
      \thead[l]{10 \textit{RPS13} intron}                        & \thead[l]{C. albicans}                               & \thead[l]{chr6:115411-115298}                         & \thead[r]{114}      & \thead[l]{RPS13: 105929-103938} \\
      \hline
      \thead[l]{11 lysyl-tRNA synthetase intron / 5$^\prime$ UTR} & \thead[l]{A. fumigatus}                              & \thead[l]{chr1:1270323-1270266}                       & \thead[r]{58}       & \thead[l]{AFUA\_1G04460: 1270442-1268328} \\
      \hline
      \thead[l]{12 metacaspase 1 intron}                         & \thead[l]{\textit{N. crassa}}                        & \thead[l]{chr7:2633707-2633620}                       & \thead[r]{88}       & \thead[l]{NCU02400: 2635820-2631309} \\
      \hline
      \thead[l]{13 thioredoxin 3$^\prime$ UTR}                    & \thead[l]{\textit{A. fumigatus}\\\textit{N. crassa}} & \thead[l]{chr4:2371534-2371435\\chr1:2599913-2600026} & \thead[r]{100\\114} & \thead[l]{AFUA\_4G09080: 2360133-2358123\\NCU02520: 2598003-2599740}  \\
      \hline
      \thead[l]{14 RING finger domain protein 3$^\prime$ UTR}     & \thead[l]{\textit{A. fumigatus}\\\textit{N. crassa}} & \thead[l]{chr7:1436607-1436525\\chr2:2021057-2021142} & \thead[r]{83\\86}   & \thead[l]{AFUA\_7G05850: 1435276-1433710\\NCU06763: 2019543-2021559}  \\
      \hline
      \thead[l]{15 thioesterase family protein 3$^\prime$ UTR}    & \thead[l]{\textit{A. fumigatus}}                     & \thead[l]{chr2:1904008-1904176}                       & \thead[r]{169}      & \thead[l]{AFUA\_2G07440: 1903479-1903916} \\
      \hline
      \thead[l]{16 \textit{GLY1} 3$^\prime$ UTR}                  & \thead[l]{\textit{S. cerevisiae}}                    & \thead[l]{chr5:67409-67247}                           & \thead[r]{163}      & \thead[l]{GLY1: 68792-67629} \\
      \hline
      \thead[l]{17 {\textit{MET13} 3$^\prime$ UTR}}               & \thead[l]{\textit{C. albicans}}                      & \thead[l]{chr3:627997-627954}                         & \thead[r]{44}       & \thead[l]{MET13: 629936-628092} \\
      \hline
      \hline
    \end{tabular}
    \captionsetup{width=25cm}
    \caption*{Table S2. \textbf{Details of the 17 fungal structural RNAs.} The NCBI accessions for the first chromosome of the five query alignments are the following: \textit{S. cerevisiae} (BK006935.2), \textit{C. albicans} (CP017623.1), \textit{N. crassa} (NC\_026501.1), \textit{A. fumigatus} (NC\_007194.1), and \textit{S. pombe} (CU329670.1).}
  \end{center}
  \end{table}
\end{landscape}

\pagebreak
\begin{table}[H]

    Table S3. \textbf{106 hits removed across the five query genomes.}
  \bigbreak
  
  \footnotesize
  \begin{tabular}{|l|l|c|c|l|}
    \hline
    \thead[c]{\textbf{genome}}                            &
    \thead[c]{\textbf{Intergenic region}\\chr:start-end}  &
    \thead[c]{\textbf{Alignment}\\avg seq len}            &
    \thead[c]{\textbf{Rfam hit}}                          &
    \thead[c]{\textbf{Rfam description}}\\
    \hline
    \textit{A. fumigatus} & 1:913875-917095 & 217 & RF00001 & 5S rRNA\\
    \textit{A. fumigatus} & 1:966944-968820 & 223 & RF00001 & 5S rRNA\\
    \textit{A. fumigatus} & 1:986548-987545 & 233 & RF01269 & Small nucleolar RNA snR80\\
    \textit{A. fumigatus} & 1:1387321-1388627 & 698 & RF00009 & Nuclear RNase P\\
    \textit{A. fumigatus} & 1:1991130-1991861 & 330 & RF00003 & U1 spliceosomal RNA\\
    \textit{A. fumigatus} & 2:4172233-4172658 & 126 & RF00001 & 5S rRNA\\
    \textit{A. fumigatus} & 2:4534354-4534839 & 241 & RF01261 & Small nucleolar RNA snR82\\
    \textit{A. fumigatus} & 3:638227-639225 & 203 & RF00001 & 5S rRNA\\
    \textit{A. fumigatus} & 3:687949-688947 & 200 & RF00001 & 5S rRNA\\
    \textit{A. fumigatus} & 3:1089131-1090129 & 231 & RF00001 & 5S rRNA\\
    \textit{A. fumigatus} & 3:3323928-3325304 & 183 & RF00005 & tRNA\\
    \textit{A. fumigatus} & 3:3785055-3785278 & 196 & RF01503 & A. fumigatus sRNA Afu 203\\
    \textit{A. fumigatus} & 4:755217-756213 & 186 & RF00001 & 5S rRNA\\
    \textit{A. fumigatus} & 4:757715-758711 & 240 & RF00001 & 5S rRNA\\
    \textit{A. fumigatus} & 4:1996559-1997556 & 282 & RF01496 & A. fumigatus sRNA Afu 182\\
    \textit{A. fumigatus} & 4:3162957-3163743 & 188 & RF00001 & 5S rRNA\\
    \textit{A. fumigatus} & 4:3279339-3280517 & 341 & RF00003 & U1 spliceosomal RNA\\
    \textit{A. fumigatus} & 5:309691-310445 & 131 & RF00005 & tRNA\\
    \textit{A. fumigatus} & 5:2496656-2497518 & 539 & RF01434 & Small nucleolar RNA snR3\\
    \textit{A. fumigatus} & 5:2631624-2636010 & 182 & RF02543 & Eukaryotic LSU rRNA\\
    \textit{A. fumigatus} & 6:31672-32555 & 160 & RF02543 & Eukaryotic LSU rRNA\\
    \textit{A. fumigatus} & 6:1654992-1656125 & 430 & RF00005 & tRNA\\
    \textit{A. fumigatus} & 6:1656782-1660223 & 495 & RF00005 & tRNA\\
    \textit{A. fumigatus} & 6:2787410-2788874 & 462 & RF01239 & Small nucleolar RNA snR49\\
    \textit{A. fumigatus} & 6:3000535-3001451 & 244 & RF00001 & 5S rRNA\\
    \textit{A. fumigatus} & 6:3477308-3478208 & 183 & RF00005 & tRNA\\
    \textit{A. fumigatus} & 7:319031-319850 & 251 & RF01240 & Small nucleolar RNA snR85\\
    \textit{A. fumigatus} & 7:662854-664651 & 342 & RF01445 & Small nucleolar RNA snR94\\
    \textit{A. fumigatus} & 7:817049-817258 & 118 & RF00005 & tRNA\\
    \textit{A. fumigatus} & 8:422221-423216 & 233 & RF00001 & 5S rRNA\\
    \textit{A. fumigatus} & 8:1542740-1545898 & 133 & RF00005 & tRNA\\
    \textit{A. fumigatus} & 8:1792689-1793274 & 113 & RF00001 & 5S rRNA\\
    \hline
  \end{tabular}


  \bigbreak
  
  \begin{tabular}{|l|l|c|c|l|}
    \hline
    \thead[c]{\textbf{genome}}                            &
    \thead[c]{\textbf{Intergenic region}\\chr:start-end}  &
    \thead[c]{\textbf{Alignment}\\avg seq len}            &
    \thead[c]{\textbf{Rfam hit}}                          &
    \thead[c]{\textbf{Rfam description}}\\
    \hline
    \textit{C. albicans} & 1:1095960-1097142 & 167 & RF01245 & Small nucleolar RNA snR9\\
    \textit{C. albicans} & 1:1862322-1862510 & 120 & RF00005 & tRNA\\
    \textit{C. albicans} & 1:2384577-2386237 & 219 & RF00005 & tRNA\\
    \textit{C. albicans} & 1:2587936-2588554 & 143 & RF00005 & tRNA\\
    \textit{C. albicans} & 1:2960366-2960890 & 204 & RF01258 & Small nucleolar RNA snR10\\
    \textit{C. albicans} & 1:3120122-3120929 & 359 & RF01267 & Small nucleolar RNA snR37\\
    \textit{C. albicans} & 2:638517-639334 & 136 & RF00005 & tRNA\\
    \textit{C. albicans} & 2:833442-833939 & 232 & RF01260 & Small nucleolar RNA snR11\\
    \textit{C. albicans} & 2:1077440-1077328 & 127 & RF00005 & tRNA\\
    \textit{C. albicans} & 2:1497591-1498446 & 244 & RF01266 & Small nucleolar RNA snR45\\
    \textit{C. albicans} & 2:1620125-1620260 & 115 & RF00005 & tRNA\\
    \textit{C. albicans} & 2:2179215-2179654 & 263 & RF01256 & Small nucleolar RNA snR43\\
    \textit{C. albicans} & 3:458225-459259 & 351 & RF01269 & Small nucleolar RNA snR80\\
    \textit{C. albicans} & 3:681093-681277 & 155 & RF00005 & tRNA\\
    \textit{C. albicans} & 3:1734285-1734632 & 264 & RF01243 & Small nucleolar RNA snR33\\
    \textit{C. albicans} & 4:902632-903051 & 118 & RF00005 & tRNA\\
    \textit{C. albicans} & 4:1251386-1251607 & 172 & RF00005 & tRNA\\
    \textit{C. albicans} & 4:1324975-1325487 & 123 & RF00005 & tRNA\\
    \textit{C. albicans} & 4:1381379-1382068 & 213 & RF01242 & Small nucleolar RNA snR36\\
    \textit{C. albicans} & 5:201169-201945 & 157 & RF00005 & tRNA\\
    \textit{C. albicans} & 5:305503-306587 & 170 & RF01960 & Eukaryotic SSU rRNA\\
    \textit{C. albicans} & 5:430140-430754 & 133 & RF00005 & tRNA\\
    \textit{C. albicans} & 6:171333-172240 & 267 & RF00004 & U2 spliceosomal RNA\\
    \textit{C. albicans} & 7:839560-840835 & 171 & RF00005 & tRNA\\
    \textit{C. albicans} & R:807594-808432 & 152 & RF01253 & Small nucleolar RNA snR46\\
    \textit{C. albicans} & R:1164444-1164566 & 112 & RF01255 & Small nucleolar RNA snR35\\
    \textit{C. albicans} & R:1830002-1831039 & 230 & RF01254 & Small nucleolar RNA snR34\\
    \textit{C. albicans} & R:1889053-1891234 & 557 & RF01960 & Eukaryotic SSU rRNA\\
    \textit{C. albicans} & R:1909314-1910064 & 111 & RF00005 & tRNA\\
    \hline
  \end{tabular}
\end{table}

\begin{table}

  \footnotesize
  \begin{tabular}{|l|l|c|c|l|}
    \hline
    \thead[c]{\textbf{genome}}                            &
    \thead[c]{\textbf{Intergenic region}\\chr:start-end}  &
    \thead[c]{\textbf{Alignment}\\avg seq len}            &
    \thead[c]{\textbf{Rfam hit}}                          &
    \thead[c]{\textbf{Rfam description}}\\
   \hline
   \textit{N. crassa} & 1:6442-9168 & 325 & RF00005 & tRNA\\
   \textit{N. crassa} & 1:238577-239090 & 172 & RF01247 & Small nucleolar RNA snR32\\
   \textit{N. crassa} & 1:1363798-1364791 & 122 & RF00001 & 5S rRNA\\
   \textit{N. crassa} & 1:2887936-2888376 & 192 & RF00009 & Nuclear RNase P\\
   \textit{N. crassa} & 1:2907344-2907539 & 178 & RF01513 & A. fumigatus snoRNA Afu 335\\
   \textit{N. crassa} & 1:2977600-2980995 & 143 & RF00001 & 5S rRNA\\
   \textit{N. crassa} & 1:2993018-2994253 & 129 & RF00093 & Small nucleolar RNA SNORD18\\
   \textit{N. crassa} & 1:5844823-5845352 & 101 & RF00001 & 5S rRNA\\
   \textit{N. crassa} & 2:2206339-2206812 & 122 & RF00001 & 5S rRNA\\
   \textit{N. crassa} & 2:2948495-2949193 & 482 & RF02462 & Ascomycota telomerase RNA\\
   \textit{N. crassa} & 2:3211687-3214126 & 233 & RF01512 & A. fumigatus sRNA Afu 309\\
   \textit{N. crassa} & 2:3540815-3541813 & 197 & RF00001 & 5S rRNA\\
   \textit{N. crassa} & 3:3354646-3355643 & 218 & RF01444 & Small nucleolar RNA snR92\\
   \textit{N. crassa} & 3:3728570-3729379 & 146 & RF00001 & 5S rRNA\\
   \textit{N. crassa} & 3:3981904-3982863 & 170 & RF00005 & tRNA\\
   \textit{N. crassa} & 3:3983902-3985130 & 278 & RF01242 & Small nucleolar RNA snR36\\
   \textit{N. crassa} & 3:4357563-4358022 & 224 & RF01261 & Small nucleolar RNA snR82\\
   \textit{N. crassa} & 4:555599-558380 & 358 & RF00005 & tRNA\\
   \textit{N. crassa} & 4:3609675-3610402 & 177 & RF00001 & 5S rRNA\\
   \textit{N. crassa} & 4:3928462-3929454 & 195 & RF00001 & 5S rRNA\\
   \textit{N. crassa} & 5:90084-91079 & 162 & RF02543 & Eukaryotic LSU rRNA\\
   \textit{N. crassa} & 5:416602-419043 & 220 & RF00005 & tRNA\\
   \textit{N. crassa} & 5:778325-780046 & 119 & RF00005 & tRNA\\
   \textit{N. crassa} & 5:837020-838658 & 287 & RF01434 & Small nucleolar RNA snR3\\
   \textit{N. crassa} & 5:2422699-2422964 & 122 & RF00001 & 5S rRNA\\
   \textit{N. crassa} & 5:5917842-5918648 & 316 & RF00003 & U1 spliceosomal RNA\\
   \textit{N. crassa} & 6:323571-324398 & 310 & RF00003 & U1 spliceosomal RNA\\
   \textit{N. crassa} & 6:1113628-1114624 & 276 & RF00003 & U1 spliceosomal RNA\\
   \textit{N. crassa} & 7:987119-988115 & 308 & RF00003 & U1 spliceosomal RNA\\
   \textit{N. crassa} & 7:3654557-3655034 & 225 & RF01248 & Small nucleolar RNA snR8\\
   \textit{N. crassa} & 7:4139782-4140779 & 194 & RF00001 & 5S rRNA\\
   \hline
  \end{tabular}

  \bigbreak
  \footnotesize
  \begin{tabular}{|l|l|c|c|l|}
    \hline
    \thead[c]{\textbf{genome}}                            &
    \thead[c]{\textbf{Intergenic region}\\chr:start-end}  &
    \thead[c]{\textbf{Alignment}\\avg seq len}            &
    \thead[c]{\textbf{Rfam hit}}                          &
    \thead[c]{\textbf{Rfam description}}\\
   \hline
   \textit{S. cerevisiae} & 4:158735-159601 & 196 & RF02543 & Eukaryotic LSU rRNA\\
   \textit{S. cerevisiae} & 7:2079-2790 & 147 & RF00005 & tRNA\\
   \textit{S. cerevisiae} & 12:451963-452438 & 401 & RF02543 & Eukaryotic LSU rRNA\\
   \textit{S. cerevisiae} & 12:453131-454122 & 802 & RF02543 & Eukaryotic LSU rRNA\\
   \textit{S. cerevisiae} & 12:454534-455180 & 567 & RF02543 & Eukaryotic LSU rRNA\\
    \hline
  \end{tabular}

    \bigbreak
  
  \begin{tabular}{|l|l|c|c|l|}
    \hline
    \thead[c]{\textbf{genome}}                            &
    \thead[c]{\textbf{Intergenic region}\\chr:start-end}  &
    \thead[c]{\textbf{Alignment}\\avg seq len}            &
    \thead[c]{\textbf{Rfam hit}}                          &
    \thead[c]{\textbf{Rfam description}}\\
    \hline
    \textit{S. pombe} & 1:952610-952873 & 143 & RF00005 & tRNA\\
    \textit{S. pombe} & 1:1746469-1746913 & 127 & RF00005 & tRNA\\
    \textit{S. pombe} & 1:1845709-1846109 & 178 & RF00001 & 5S rRNA\\
    \textit{S. pombe} & 1:3157303-3157492 & 108 & RF00005 & tRNA\\
    \textit{S. pombe} & 2:323950-324167 & 124 & RF00005 & tRNA\\
    \textit{S. pombe} & 2:1602266-1602771 & 162 & RF00005 & tRNA\\
    \textit{S. pombe} & 2:2322721-2323014 & 177 & RF00001 & 5S rRNA\\
    \textit{S. pombe} & 2:2948203-2948644 & 167 & RF00001 & 5S rRNA\\
    \textit{S. pombe} & 3:814711-815084 & 284 & RF02543 & Eukaryotic LSU rRNA\\
    \hline
  \end{tabular}
  \caption*{Table S3.  Structures that satisfied the selection criteria but had a significant hit to any Rfam families were removed. The list of 106 removed hits includes: 35 rRNAs, 32 tRNAs, 25 snoRNAs, and 15 other RNA genes.}
\end{table}


\renewcommand{\thefigure}{S\arabic{figure}}
\renewcommand{\thetable}{S\arabic{figure}}

\setcounter{figure}{0}
\section{Supplemental Figures}
\begin{figure}[!htb]
	\begin{subfigure}{1\textwidth}
		\center
		\includegraphics[width=7in, height=7in,keepaspectratio]{figures/figure_S1.pdf}	
	\end{subfigure}
	
	\caption[Results of optimized method in positive controls]{\textbf{Results of optimized method in positive controls}. When optimizing our method for discovering structural non-coding RNAs, we tested how two parameters, the number of iterative homology searches and E-value threshold, affected the number of statistically significant covariations (inferred as conserved base pairs) detected in alignments of these known functional ncRNAs. This set of positive controls included 80 structural ncRNA genes from the \textit{S. cerevisiae} genome, including all 29 H/ACA box small nucleolar RNAs (snoRNA), 5S and 5.8S ribosomal RNAs (rRNA), 39 transfer RNAs (tRNA) with one per anticodon, 5 spliceosomal RNAs (snRNA), and 5 other RNAs (RNase MRP, RNase P, SRP RNA, Telomerase RNA, and U3).  These sequences were flanked with upstream and downstream intergenic sequences, to simulate a screen for standalone ncRNA genes using the unflanked mode. With the optimized unflanked mode (see Main Fig. 2), 62 out of 80 (78\%) ncRNAs have 3 or more conserved base pairs, and only 10 (13\%) have fewer than 3 conserved base pairs when more than 25\% of base pairs are expected to covary (measure of covariation power). 8 (10\%) of these ncRNAs have less than a 25\% of base pairs expected to covary, and in all cases, these ncRNAs have low covariation power because homology is only detected in the \textit{Saccharomyces sensu stricto}. In some cases, fewer conserved base pairs are found than expected because nhmmer does not produce alignments that account for secondary structure or because of peculiarities in the query sequence. For example, the U2 sequence from \textit{S. cerevisiae} is unusually large compared to homologs from other fungi (1). Using a sequence more representative of fungal U2s such as the one from \textit{S. pombe} resulted in detection of many more conserved base pairs. All ncRNAs except H/ACA box snoRNAs and tRNAs are labeled on the plot.}
	\label{fig:S1}
\end{figure}

\begin{figure}
	\begin{subfigure}{1\textwidth}
		\center
		\includegraphics[width=6.5in, height=8in,keepaspectratio]{figures/figure_S2.pdf}	
	\end{subfigure}
	
	\caption[Example of a genomic region screened]{\textbf{Example of a genomic region screened}. Here, a 6 kb region of the \textit{S. cerevisiae} genome is shown with annotations from the most recent (01/13/2015) annotation from the \textit{Saccharomyces} Genome Database. The annotations give the coordinates of protein-coding genes, non-coding genes, and other features like transposable elements. The coordinates of introns are also provided. Based on this annotation, we defined intergenic regions (IGRs) as any unannotated regions. If intronic coordinates are explicitly provided (as for \textit{S. cerevisiae} and \textit{S. pombe}), we use those coordinates. In the annotations of the other three query genomes, the intron coordinates are not explicitly defined, so we define them as the intervening sequence between two annotated exons of the same protein-coding gene. If the coordinates of untranslated regions (UTRs) are provided (only for \textit{S. pombe}), then we use those coordinates. In the four other genomes, the IGR coordinates were treated as the UTR coordinates, with the direction of transcription of the adjacent gene used to designate the UTR as either 5$^\prime$ or 3$^\prime$.}
	\label{fig:S4}
\end{figure}

\newpage
\subsection{Experiments with positive control guanine riboswitch}
\begin{figure}[h]
	\begin{subfigure}{1\textwidth}
		\center
		\includegraphics[width=6.5in, height=8in,keepaspectratio]{figures/figure_S3.pdf}	
	\end{subfigure}
	
	\caption[Experiments with guanine riboswitch]{\textbf{Experiments with guanine riboswitch}. \textbf{(a)} The sequence and structure of the \textit{B. subtilis} \textit{xpt-pbuX} guanine riboswitch. Guanine residues are bolded. The first three guanine residues (in asterisks) were added to the beginning of the sequence to increase efficiency of in vitro transcription. \textbf{(b)} In-line probing of guanine riboswitch: red asterisks indicate the regions of differential degradation between the lane incubated with 10 $\mu$M guanine (+guanine) and those incubated with no ligand (-) or 10 $\mu$M adenine (+adenine). The no reaction (NR) lane corresponds to undigested, precursor RNA, the T1 lane corresponds to the RNA digested with RNase T1 which cleaves after guanine residues, and -OH lane corresponds to the RNA under partial alkaline digestion, which produces a single nucleotide ladder. Typically, structured regions are less susceptible to degradation, and our gel is concordant with this: the base pairing regions labeled ``P1" to ``P3" are less frequently the site of degradation than unpaired regions. The regions of differential degradation are also consistent with gels from the literature (2). \textbf{(c)} Representative isothermal titration calorimetry results for the guanine riboswitch. The top window in each plot shows the raw ITC data and the bottom window shows $\Delta$H. 1.5 $\mu$M RNA was titrated against 15 $\mu$M guanine (left) or adenine (right). Only titration with guanine results in the expected sigmoidal binding isotherm and has derived thermodynamic parameters consistent with the literature (3).}
	\label{fig:S2}
\end{figure}

\newpage
\subsection{Experiments with \textit{GLY1} 3$^\prime$ UTR and \textit{MET13} 3$^\prime$ UTR}
\begin{figure}[h]
	\begin{subfigure}{1\textwidth}
		\center
		\includegraphics[width=6.5in, height=8in,keepaspectratio]{figures/figure_S4.pdf}	
	\end{subfigure}
	
	\caption[Experiments with the \textit{GLY1} 3$^\prime$ UTR structure]{\textbf{Experiments with the \textit{GLY1} 3$^\prime$ UTR structure}. \textbf{(a)} The sequence and proposed structure of the \textit{GLY1} 3$^\prime$ UTR structure from \textit{S. cerevisiae}. Guanine residues are bolded. The first three guanine residues (in asterisks) were added to the beginning of the sequence to increase efficiency of in vitro transcription. \textbf{(b)} Two gels from the in-line probing assay are shown. In each case, half of the sample was run initially at 50 W on a sequencing gel. After three hours, the second half was loaded. Therefore, the first half was run for 6 hours total to increase resolution of the 3$^\prime$ end of the RNA. There is no differential degradation between the RNA incubated with no ligand (-) compared to those incubated with 10 mM glycine, 10 mM L-threonine, or a mixture of several other compounds (L-serine, tetrahydrofolate, L-aspartate, guanine, adenine, PLP, and TPP) all at 10 mM. The no reaction (NR) lane corresponds to undigested, precursor RNA, the T1 lane corresponds to the RNA digested with RNase T1 which cleaves after guanine residues, and the -OH lane corresponds to the RNA under partial alkaline digestion, which produces a single nucleotide ladder. Typically, structured regions are less susceptible to degradation, and our gel is concordant with this: the base pairing regions labeled ``P1" to ``P4" are less frequently the site of degradation than unpaired regions. \textbf{(c)} Representative isothermal titration calorimetry results. The top window in each plot shows the raw ITC data and the bottom window shows $\Delta$H. 5 $\mu$M RNA was titrated against 100 $\mu$M of either glycine, threonine, or the mixture of other ligands. None resulted in the sigmoidal binding isotherm expected if a ligand were binding to the RNA.}
	\label{fig:S3}
\end{figure}

\begin{figure}
	\begin{subfigure}{1\textwidth}
		\center
		\includegraphics[width=6.5in, height=8in,keepaspectratio]{figures/figure_S5.pdf}	
	\end{subfigure}
	
	\caption[Experiments with the \textit{MET13} 3$^\prime$ UTR structure]{\textbf{Experiments with the \textit{MET13} 3$^\prime$ UTR structure}. \textbf{(a)} The sequence and proposed structure of the \textit{MET13} 3$^\prime$ UTR structure from \textit{H. burtonii}. Guanine residues are bolded. The first three guanine residues (in asterisks) were added to the beginning of the sequence to increase efficiency of in vitro transcription. \textbf{(b)} Results from the in-line probing assay are shown. There is no differential degradation between the RNA incubated with no ligand (-) compared to those incubated with 250 $\mu$M SAM or SAH. The no reaction (NR) lane corresponds to undigested, precursor RNA, the T1 lane corresponds to the RNA digested with RNase T1 which cleaves after guanine residues, and the -OH lane corresponds to the RNA under partial alkaline digestion, which produces a single nucleotide ladder. \textbf{(c)} Representative isothermal titration calorimetry results. The top window in each plot shows the raw ITC data and the bottom window shows $\Delta$H. On the left, 12 $\mu$M RNA was titrated against 150 $\mu$M SAM and on the right 8 $\mu$M RNA was titrated against 180 $\mu$M SAH. Neither resulted in the sigmoidal binding isotherm expected if a ligand were binding to the RNA.}
	\label{fig:S4}
\end{figure}

\newpage

\section{Supplemental Information}

\subsection{Fungal genomes include in database}

A list in alphabetical order of the 1371 Ascomycota fungal genomes in the whole genome database used for homology search is included. 

\subsection{High-quality genome annotations used to determine if structures are associated with mRNAs}

To determine whether a structural ncRNA candidate was located within an UTR or intron of mRNA, we examined if its representatives were consistently located adjacent or within the annotated mRNA of that gene, respectively. The following 24 relatively complete annotations from Genbank were used for this purpose. For all candidate structures, the genomic coordinates of any representatives from a genome in this list were used to determine its genomic context.

\textit{Ascodesmis nigricans}, \textit{Aspergillus fumigatus}, \textit{Aspergillus oryzae}, \textit{Candida albicans}, \textit{Candida viswanathii}, \textit{Debaryomyces hansenii}, \textit{Eremothecium gossypii}, \textit{Fusarium verticillioides},
\textit{Kazachstania africana}, \textit{Kluyveromyces lactis}, 
\textit{Lachancea thermotolerans}, \textit{Naumovozyma dairenensis},
\textit{Neosartorya fischeri}, \textit{Neurospora crassa}, \textit{Penicillium chrysogenum}, \textit{Purpureocillium lilacinum}, \textit{Rasamsonia emersonii}, \textit{Saccharomyces cerevisiae}, \textit{Scheffersomyces stipitis}, \textit{Schizosaccharomyces pombe},
\textit{Talaromyces stipitatus}, \textit{Thermothelomyces thermophilus}, 
\textit{Torrubiella hemipterigena}, and \textit{Uncinocarpus reesii}.

Structures we have classified as within UTRs or introns are consistent with those locations in all of the above genomes that have homologs. For example, structure 11 is found in 7 of these genomes. We designated it as associated with the lysyl-tRNA synthetase because it is found in an intron of that gene in \textit{A. fumigatus}, \textit{A. oryzae}, \textit{P. chrysogenum}, \textit{T. stipitatus}, and \textit{T. hemipterigena}, and in its 5$^\prime$ UTR in \textit{N. fischeri} and \textit{P. lilacinum}. 

\subsection{Alignments of structural ncRNA candidates}

The alignments of the 17 structural ncRNAs are available in a zip folder. The alignments are in Stockholm format, the file format used by all software included in our method (HMMER, Infernal, and R-scape). 

In some cases, running R-scape (v1.5.16) with the -s --fold options will produce structures that appear slightly different than those presented in the main figures. In most cases, this is because we individually trimmed the R2R renderings of some structures to include only the regions with conserved structure.

For H/ACA structures 2 and 3, the alignment resulting from our method produced many possibly spurious covariations, likely because the sequences were initially aligned using nhmmer, which generates an alignment based only on primary sequence conservation. This issue is described in more detail in the section ``Misalignments can induce spurious covariations" from the preprint ``Evolutionary conservation of RNA sequence and structure," available at \url{http://rivaslab.org/publications.html}.  For those two cases only, a covariance model was built using the proposed structure after iterative nhmmer homology search and a single sequence from a model organism (\textit{C. albicans} for structure 2 and \textit{A. fumigatus} for structure 3). This covariance model was calibrated and used to search against the hits from the initial alignment, with an E-value of $10^{-5}$, to produce a more structurally-informed alignment. R-scape -s --fold was then run on this alignment and displayed in the figures shown here.

\subsection{Sequences used in experiments}

\subsubsection{For guanine riboswitch}
Before using in-line probing and isothermal titration calorimetry on our experimental RNAs, we first performed these assays on a positive control, the guanine riboswitch. 

The following sequence from the \textit{B. subtilis} \textit{xpt-pbuX} guanine riboswitch was placed in pUC19 plasmid between the BamHI and EcoRI restrictions downstream of the T7 RNA Polymerase promoter sequence (5$^\prime$-TAATACGACTCACTATAG-3$^\prime$). Three Gs were added directly after the promoter sequence to enhance efficiency of transcription.

5$^\prime$-AATATAATAGGAACACTCATATAATCGCGTGGATATGGCACGCAAGTTTCTA\\CCGGGCACCGTAAATGTCCGACTATGGGTGAGCAATGGAAC-3$^\prime$

Prior to in vitro transcription, PCR amplification of the plasmid was performed using the following primers: a forward primer (5$^\prime$-GCTATGACCATGATTACG-3$^\prime$) that anneals upstream in the pUC19 backbone and a reverse primer (5$^\prime$-GTTCCATTGCTCACCCATAG-3$^\prime$) that is the reverse complement of the end of the inserted sequence. 

\subsubsection{For the \textit{GLY1} 3$^\prime$ UTR structure}

The following sequence corresponding to the \textit{GLY1} 3$^\prime$ UTR structure from \textit{S. cerevisiae} was placed in pUC19 plasmid between the BamHI and EcoRI restriction sites downstream of the T7 RNA Polymerase promoter sequence. Three Gs were added directly after the promoter sequence to enhance efficiency of transcription.

5$^\prime$-
CACATTTGAGAACCAATGGTTAGTTTCCAGCGCACCTAGCGACTAACTACCCTA\\
TTATGGAGGGACGAAGGTGGTCTTTCTGAGGGAAGGAGGAAAAAAAGGTAAGAACCT\\
CTTTTCTTTGTTCCCCATTGAGAGATCGACTGAATATGTTTGGTTTTTTTGG-3$^\prime$

Prior to in vitro transcription, PCR amplification of the plasmid was performed using the following primers: a forward primer (5$^\prime$-GCTATGACCATGATTACG-3$^\prime$) that anneals upstream in the pUC19 backbone and a reverse primer (5$^\prime$-GAATTCCCAAAAAAACCAAACA-3$^\prime$) that is the reverse complement of the EcoRI restriction site (3$^\prime$-CTTAAG-5$^\prime$) and end of the inserted sequence.

\subsubsection{For the \textit{MET13} 3$^\prime$ UTR structure}

The following sequence corresponding to the \textit{MET13} 3$^\prime$ UTR structure from \textit{Hyphopichia burtonii} was synthesized as an oligonucleotide. Because the stem has only 5 base pairs, this was chosen instead of the \textit{C. albicans} sequence which contains a wobble base pair that may be less stable in vitro.

Its sequence is:
5$^\prime$-CAGGATATTTAGTGCACTCAACATGCACATCCTTAAAGTCGATTTC\\AACA-3$^\prime$. PCR amplification was performed using a forward primer (5$^\prime$-TAATACGACTCACTATA\\GGGCAGGATATTTAGTGCACTCAAC-3$^\prime$) which contains the T7 RNA Polymerase promoter, three Gs to increase transcriptional efficiency, and a 22-nucleotide region identical to the start of the desired amplified sequence. The reverse primer used was  5$^\prime$-TGTTGAAATCGACTTTAAGGATG\\TG-3$^\prime$.

\section{References}

\phantom{1234}

\noindent
1. Ares Jr, M. (1986)
U2 RNA from yeast is unexpectedly large and contains homology to vertebrate U4, U5, and U6 small nuclear RNAs.
\textit{Cell}, \textbf{47}, 49--59.

\noindent
2. Mandal, M., Boese, B., Barrick, J.E., Winkler, W.C., and Breaker, R.R. (2003)
Riboswitches control fundamental biochemical pathways in \textit{Bacillus subtilis} and other bacteria. 
\textit{Cell}, \textbf{113}, 577--586.

\noindent
3. Gilbert, S.D., Stoddard, C.D., Wise, S.J., and Batey, R.T. (2006)
Thermodynamic and kinetic characterization of ligand binding to the purine riboswitch aptamer domain.
\textit{Journal of Molecular Biology}, \textbf{359}, 754--768.

\pagestyle{plain}

\end{document}
