## Supplementary figures and images for "Discovery of 17 conserved structural RNAs in fungi"

### figure_S1.pdf

#### positive control screen

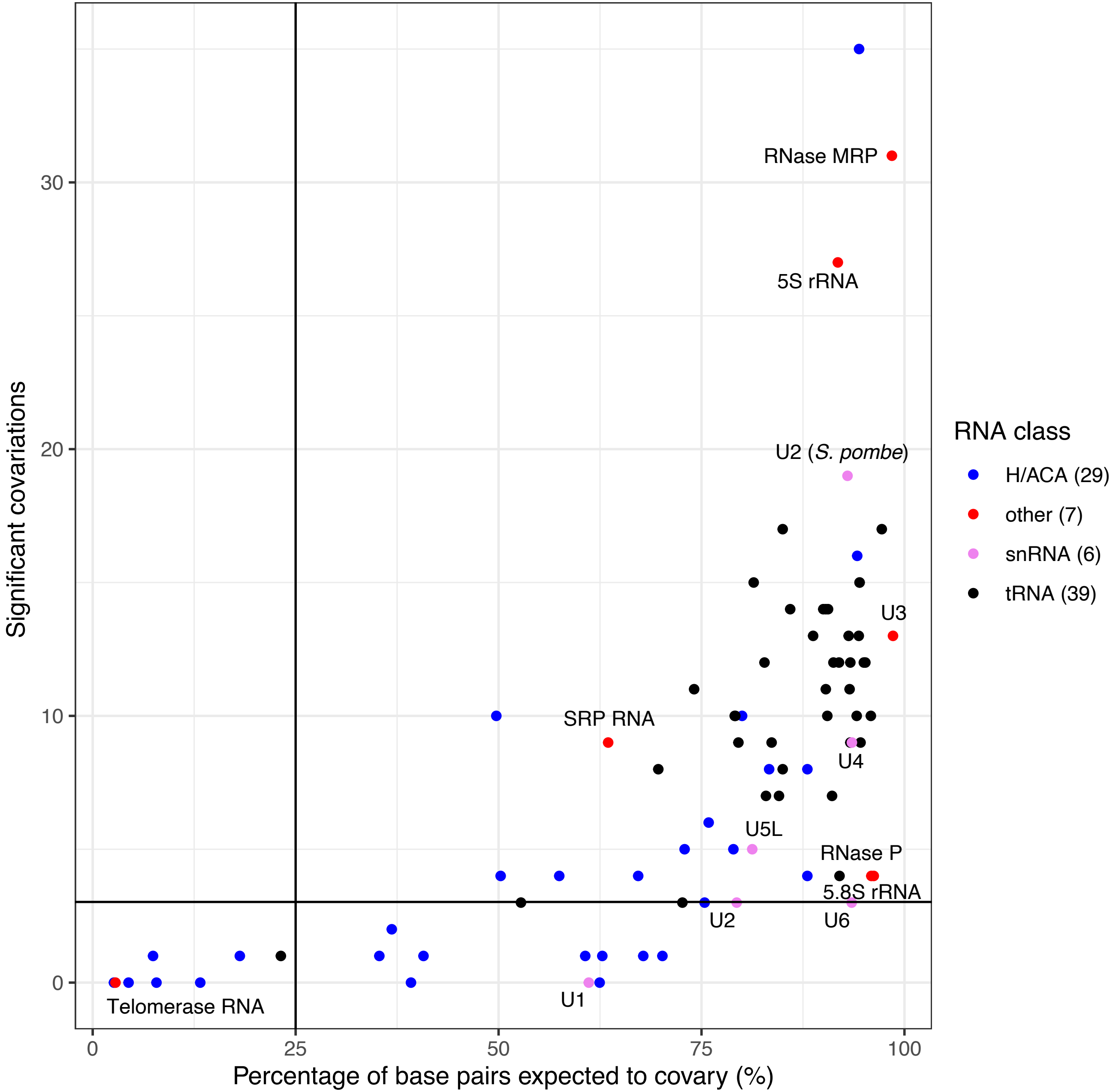

### figure_S2.pdf

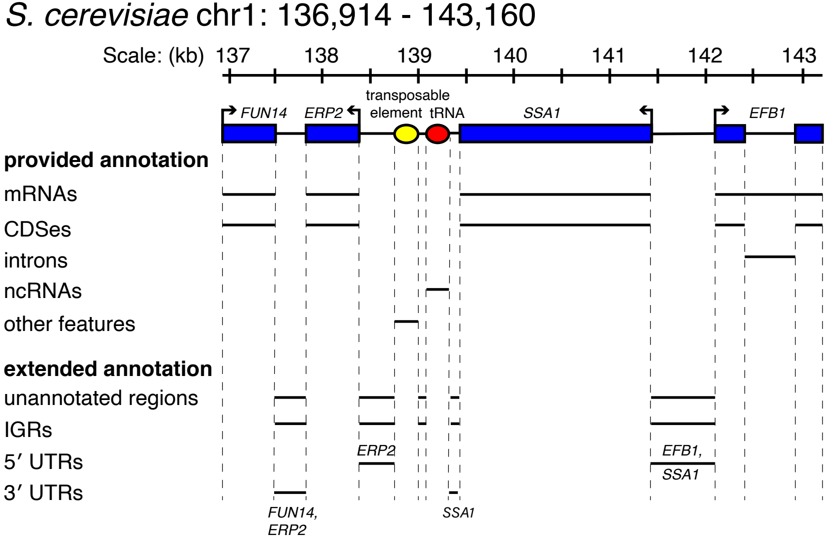

### figure_S3.pdf

guanine riboswitch (positive control)

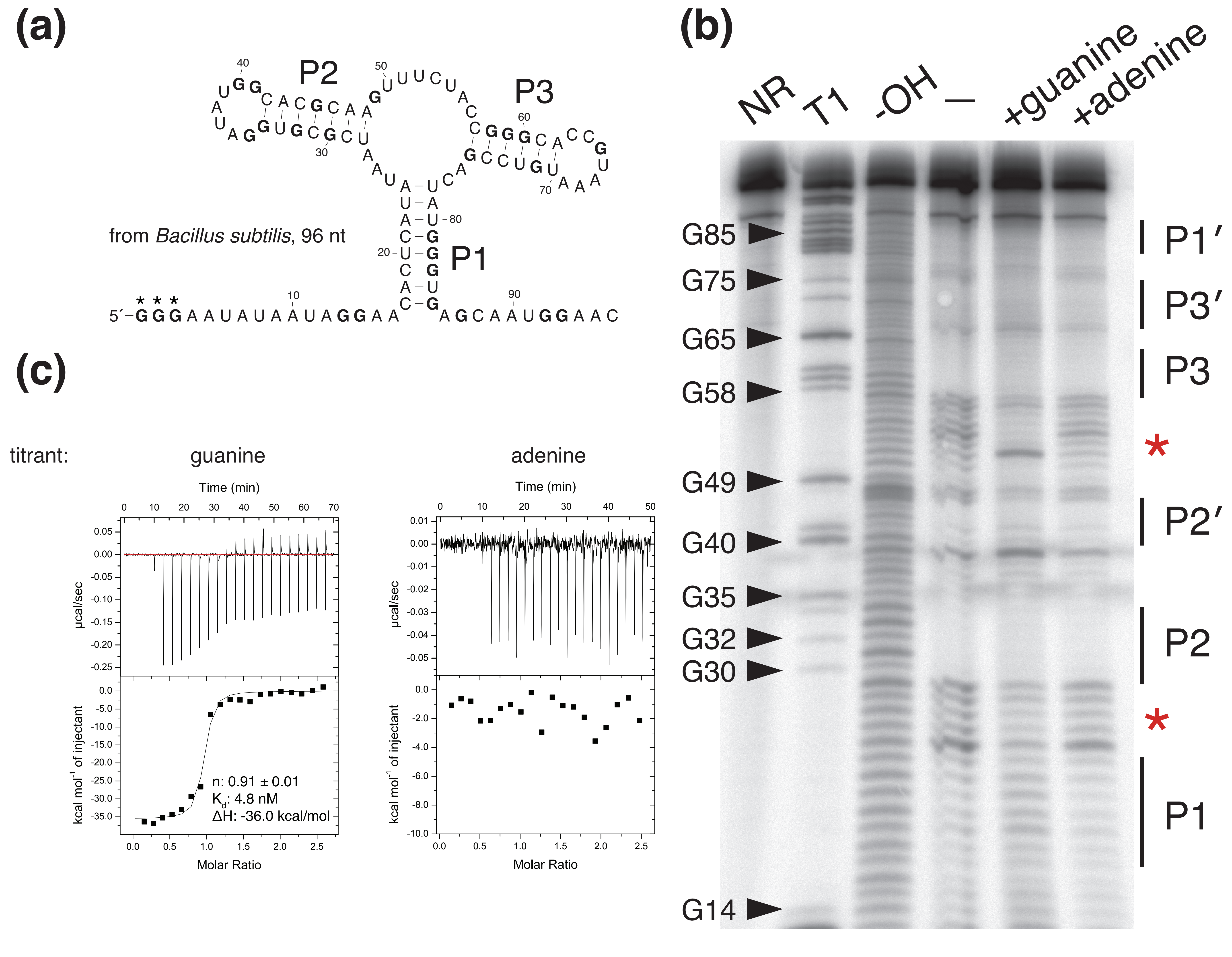

### figure_S4.pdf

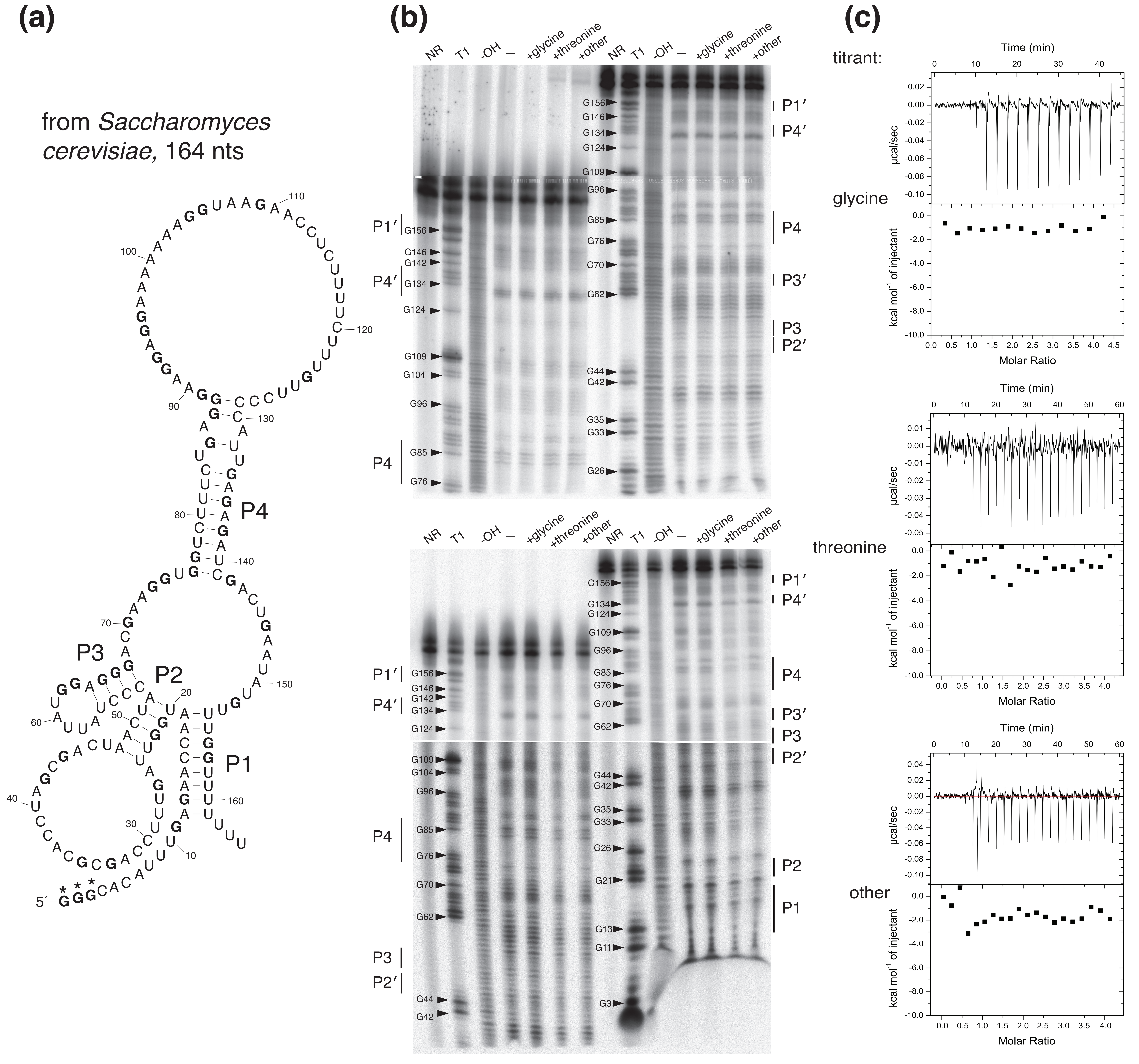

### figure_S5.pdf

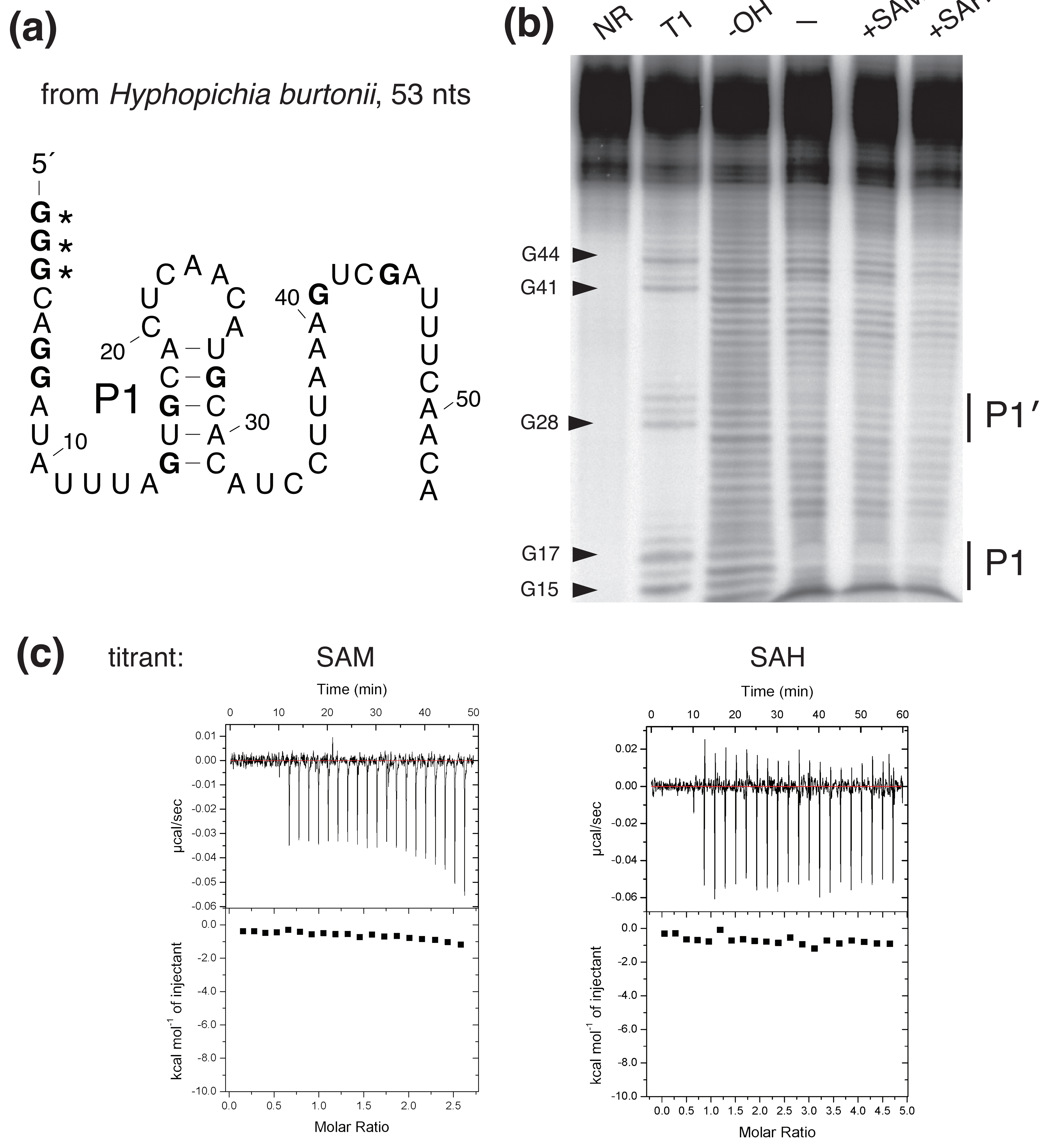
